# Using mechanistic models to highlight research priorities for tick-borne zoonotic diseases: Improving our understanding of the ecology and maintenance of Kyasanur Forest Disease in India

**DOI:** 10.1101/2022.10.17.512427

**Authors:** Richard Hassall, Sarah J. Burthe, Stefanie M. Schäfer, Nienke Hartemink, Bethan V. Purse

## Abstract

The risk of spillover of zoonotic diseases to humans is changing in response to multiple environmental and societal drivers, particularly in tropical regions where the burden of neglected zoonotic diseases is highest and land use change and forest conversion is occurring most rapidly. In these regions, neglected zoonotic diseases can have significant impacts on poor and marginalised populations in low-resource settings but ultimately receive less attention and funding for research and interventions. As such, effective control measures and interventions are often hindered by a limited ecological evidence base, which results in a limited understanding of epidemiologically relevant hosts or vectors and the processes that contribute to the maintenance of pathogens and spillover to humans. Here, we develop a generalisable next generation matrix modelling framework to better understand the transmission processes and hosts that have the greatest contribution to the maintenance of tick-borne diseases with the aim of improving the ecological evidence base and framing future research priorities for tick-borne diseases. Using this model we explore the relative contribution of different host groups and transmission routes to the maintenance of a neglected zoonotic tick-borne disease, Kyasanur Forest Disease Virus (KFD). The results highlight the potential importance of transovarial transmission and small mammals and birds in maintaining this disease. This contradicts previous hypotheses that primates play an important role influencing the distribution of infected ticks. There is also a suggestion that risk could vary across different habitat types. In light of these results we outline the key knowledge gaps for this system and future research priorities that would aid in informing effective interventions and control measures.

## Introduction

The global burden of zoonotic diseases is changing in response to multiple environmental change drivers including climate change, increased international trade and travel, and human modification of ecosystems through intensive farming, dams and irrigation, deforestation, human settlement and urbanisation (Jones et al., 2013). The mechanisms underpinning spillover of zoonotic diseases to humans can be complex and are not well understood, hampering design and implementation of effective interventions. Most zoonotic diseases have complex transmission cycles involving communities of vectors and animal hosts, with heterogeneity in disease dynamics being highly dependent on host ecology and evolutionary biology, (Karesh et al., 2012; Streicker et al., 2010; VanderWaal & Ezenwa, 2016; Webster et al., 2016). This heterogeneity in disease dynamics also needs to be considered in conjunction with human behaviour and use of ecosystems, which can govern exposure to infected vectors and hosts (Lambin et al., 2010; Murray & Daszak, 2013). As well as other social and political factors such as access to health care, land, knowledge and livelihoods, that further determine vulnerability of human communities to zoonotic diseases (Asaaga et al., 2021; Dzingirai et al., 2017; Gibb et al., 2020).

Disease control programs targeting zoonotic disease tend to utilise human-focused approaches, such as vaccination or drug treatment, with little consideration of the ecological and environmental context that results in spillover to humans (Sokolow et al., 2019; Webster et al., 2016). Effective disease control is often limited by a poor ecological evidence base, specifically by lack of knowledge of which different vector and host species, and which processes, are contributing to transmission and human spillover (Burthe et al., 2021; Webster et al., 2016). This is particularly the case for neglected zoonoses that primarily affect poor and marginalised populations in low-resource settings (Welburn et al., 2015), in which less attention and funding is available for research and interventions (Hotez & Kamath, 2009; Liese & Schubert, 2009; Webster et al., 2016). It is estimated that neglected zoonoses impact over 2 billion people causing around half a million deaths annually, though the true extent is likely several fold higher given vast under-reporting of these diseases (Herricks et al., 2017; Liese et al., 2014). New frameworks have been put forward, which aim to prevent and manage zoonotic diseases by identifying the hierarchical series of barriers that must be overcome by a pathogen to facilitate spillover from an animal reservoir into a human or livestock host (Burthe et al., 2021; Plowright et al., 2017; Sokolow et al., 2019). Given improved ecological knowledge, these barriers, such as sufficient density and competence of reservoir hosts, can be targeted by ecological interventions, complementing more conventional approaches that target humans (Burthe et al., 2021; Sokolow et al., 2019).

Exacerbating these knowledge needs for improved management, it is clear that the role of reservoir hosts and vector species, and different transmission processes, are highly sensitive to ecosystem and landscape context (Hartemink et al., 2015; Li et al., 2019). Vector and reservoir roles can not be considered as stable life history traits of species but depend on the ecosystems in which species are embedded and the relative contributions to transmission of other species within the community (M. G. Roberts & Heesterbeek, 2020). Several global change drivers, including land use change, agricultural intensification and human settlement, are hypothesised to be contributing to zoonotic disease spillover by bringing people into greater contact with domestic and wildlife animal reservoirs and vectors at the closer interfaces between human habitation, agricultural and natural habitats (Despommier et al., 2007; Plowright et al., 2017; Redding et al., 2019; Roberts et al., 2021). Furthermore, these human modified landscapes have been shown to lead to increased diversity of zoonotic hosts (Faust et al., 2017) and could result in changes in host densities and interspecies contact rates altering disease dynamics (Faust et al., 2018).

Such human-animal-environment interfaces or ecotones now dominate much of the land in tropical regions where the burden of neglected zoonotic diseases is highest and land use change and forest conversion is occurring most rapidly (Despommier et al., 2007). Within such settings, attempts to build the ecological evidence base to inform interventions must take account of potential variation in host and vector roles and their habitat associations and interactions that govern transmission processes across the human-animal-environment interface.

Models have a key role to play in predicting the variable impact of hosts and vectors on disease dynamics and infection risk across human-animal-environment interfaces and rapidly changing landscapes (Roberts et al. 2021; Webster et al. 2016) but need to be contextualised carefully to local empirical data availability and knowledge needs for the disease interventions (Asaaga et al., 2022). Models are particularly critical for understanding tick-borne disease systems (TBDs), in which ticks may feed on and transmit infection to different vertebrate hosts at different life stages via different routes of transmission. These include transovarial transmission between adult ticks and eggs, non-systemic transmission between larvae and nymphs co-feeding on the same hosts and systemic transmission between infected hosts and larval, nymph or adult ticks. Mechanistic models have been applied to understand and compare the relative contributions of transovarial, non-systemic and systemic transmission to the basic reproduction number, R_0_, between tick-borne pathogen species (Hartemink et al., 2008; Matser et al., 2009). Models have been further applied to explore the effects of vector demography, aggregation and host biodiversity on tick-borne pathogen persistence and control in different settings (Norman et al., 2016; Perkins et al., 2003; Rosà & Pugliese, 2007) though have been rarely explicitly linked to local management strategies. In their Resource-Based Habitat Concept for vector-borne diseases, (Hartemink et al., 2015) advocated for integrating functional resource use of each host, pathogen and vector species, linked to particular habitats across landscape mosaics into spatial predictive frameworks (see also (Vanwambeke et al., 2016)). Though some models have considered the impacts of variable host habitat use and tick-host interactions across space and time on transmission (Jones et al., 2011; Li et al., 2016, 2019), these are largely confined to temperate, resource-rich settings and systems such as Lyme disease and tick-borne encephalitis in the United States and Europe. For neglected zoonotic disease systems, mechanistic modelling approaches that combine available empirical data with knowledge and data from other similar disease systems, can enable insights into the key processes that may contribute to the maintenance of pathogens. Collecting empirical data can be logistically challenging and costly, and by exploring the sensitivity of models to different parameters it is possible to highlight parameters for which robust empirical estimates would reduce uncertainty in predictions whilst also identifying parameters that are less likely to be influential (Hartemink et al., 2008). This can aid in framing key knowledge gaps and potential priorities for future research and interventions.

Here, we adopt a mechanistic modelling approach for a neglected zoonotic tick-borne disease system, Kyasanur Forest Disease (KFD) in India, to gain insights into the relative contribution of different host groups (small mammals, birds and primates) and transmission routes to the maintenance of this pathogen. The latter represent priority local knowledge needs to inform improved interventions (Burthe et al., 2021).

KFD’s aetiological agent is a tick-borne flavivirus causing potentially fatal haemorrhagic disease in people in the Western Ghats region of south India, with 400-500 reported cases a year and mortality rate of up to 10% (Shah et al., 2018). Humans can contract KFD when bitten by an infected tick but are considered dead-end hosts for the disease (Murhekar et al., 2015). The disease primarily affects rural forest communities, including tribal groups and plantation and forestry workers (Mourya et al., 2013; Patil et al., 2017; Sadanandane et al., 2017). Approximately 69% of small-holder farmers and tribal groups surveyed in the region have reported that they were concerned by the impact KFD has had on their livelihoods, highlighting the impact of this disease in the region (Asaaga et al., 2020; Patil et al., 2017). Correlative modelling of recent human outbreaks indicates that the risk of virus spillover into humans is highest in diverse agro-forestry landscapes, created when moist evergreen forest is replaced with plantations and paddy cultivation (Purse et al., 2020), consistent with the hypothesis that KFD is an ecotonal disease (Pattnaik, 2006). However, we have limited knowledge of the transmission processes and hosts that are likely to contribute to the maintenance of KFD and influence the distribution of infected ticks. Thus there is an urgent requirement for understanding of the conditions and mechanisms underpinning transmission cycles in reservoirs and spillover to humans across these landscape mosaics.

KFD has a complex transmission cycle in which various tick species (mostly *Haemaphysalis* genus, but also some *Ixodes)* and vertebrate hosts, (including rodents and shrews, monkeys and birds), have been implicated (Pattnaik, 2006). Monkeys, mainly the black-footed grey langur (*Semnopithecus hypoleucos*) and the bonnet macaque (*Macaca radiata*), are hypothesised to act as amplifying hosts, by infecting large numbers of larval ticks with the virus, and are believed to create a hotspots of infections when they die (Mourya & Yadav, 2016). However, empirical evidence to support this hypothesis is lacking and there is very limited recent information on the potential role of other relevant wild hosts such as small mammals and birds following substantial land use change (Burthe et al., 2021). To date there is also a very poor understanding of the importance of transovarial transmission in the maintenance of KFD, where infection is passed from an adult female to their eggs. Although transovarial transmission has been demonstrated in laboratory studies in some native Indian Ixodes tick species such as *Ixodes petauristae (Singh et al., 1963, 1968)* there is no evidence of this occurring in *Haemaphysalis* ticks in situ (Burthe et al., 2021). If ticks are able to maintain KFD through transovarial transmission this could add significance to the potential role of cattle this system, which, while not directly acting as hosts for systemic transmission due to the fact that cattle do not develop a long-lasting viraemia (Anderson & Singh, 1971; Varma et al., 1960), may already amplify tick populations as an important blood meal host and may influence the distribution of infected ticks (Balasubramanian et al., 2019; Mehla et al., 2009). Currently, empirical knowledge of the role of different species of vector and hosts in the distribution of infected ticks and transmission cycle of KFD is lacking but management guidelines advocate for tick control on cattle and around the sites of host (monkey) deaths, despite limited empirical evidence to support these strategies (Burthe et al. 2021). Given KFD’s high human impact, sensitivity of spillover to land use change, ecotonal nature, low ecological evidence-base for interventions and complex transmission cycle, it is an ideal and important test case with which to explore the insights into host roles, transmission processes and habitat associations that can be gained by combining mechanistic models with extant empirical data and local management needs.

We utilise next generation matrix (NGM) approaches for tick-borne pathogens already developed by Hartemink et al. (2008). Previous NGM approaches investigating KFD have estimated the contribution of different modes of transmission to the R_0_ of KFD (Matser et al. 2009) but have so far considered five ‘types-at-infection’, ticks infected at different life stages (egg, larva, nymph and adult) and newly infected vertebrate hosts as a single host type. To further understand the contribution of different vertebrate hosts we build on previous models by expanding the number of host types to include small mammals, birds and primates as individual ‘types-at-infection’ as well as adapting this framework to incorporate the densities of different vertebrate hosts. This will allow us to assess the likely contribution of different host groups to systemic transmission of KFD as well as the relative contribution of transovarial transmission and non-systemic (co-feeding) transmission in maintaining KFD. In addition, we also consider the potential changes in R_0_ and reservoir host roles for KFD across different habitat types by introducing scaling factors to account for potential variation in tick abundance and subsequent burdens on hosts as a result of different cattle densities or habitat types. Using the results from these models we aim to highlight the knowledge gaps and future research priorities that would help to improve our understanding of natural foci of KFD and aid in the development of interventions.

## Methods

The NGM approach used here provides an intuitive estimate of R_0_ for tick-borne pathogens, where R_0_ is derived as the largest eigenvalue of the NGM (Diekmann et al., 1990). This eigenvalue is indicative of pathogen generations growing in size when it is greater than 1 and declining in size when the eigenvalue is less than 1 as well as providing an estimate of the per generation increase in the number of infected host or tick life stages. This approach also allows the relative contribution of different transmission routes to R_0_ to be established (Hartemink et al., 2008; Matser et al., 2009). The NGM model includes 7 types-at-infection: (1) Tick-infected-as-egg, (2) Tick-infected-as-larva, (3) Tick –infected-as-nymph, (4) Tick-infected-as-adult, (5) Newly-infected-small-mammal, (6) Newly-infected-bird, (7) Newly-infected-primate.

Given the uncertainty around transovarial transmission of KFD occurring in the wild (Burthe et al., 2021), we include a second NGM model excluding the possibility for transovarial transmission and the tick infected as egg type-at-infection. This allowed us to consider the potential impact of transovarial transmission on the persistence of KFD. Both models assume that adult ticks do not feed on primates and small mammals, based on previous data from (Rajagopalan et al., 1968) and (Trapido, Goverdhan, et al., 1964). Therefore, infected small mammals and primates can infect only larvae and nymphs (thereby producing new cases of type-at-infection 2 and 3, respectively, ticks-infected-as-larvae and ticks-infected-as-nymphs). Ticks take one blood meal per life stage, and when they get infected, they can only pass this infection on in the next life stage. This, combined with the fact that adults don’t feed on small mammals and primates, means that these hosts can only be infected by ticks that were infected as an egg (during the bloodmeal they take as larva or nymph) or by tick that were infected as a larva (during their bloodmeal as nymph). For the conceptual model see Figure 1.

**Figure 1:**
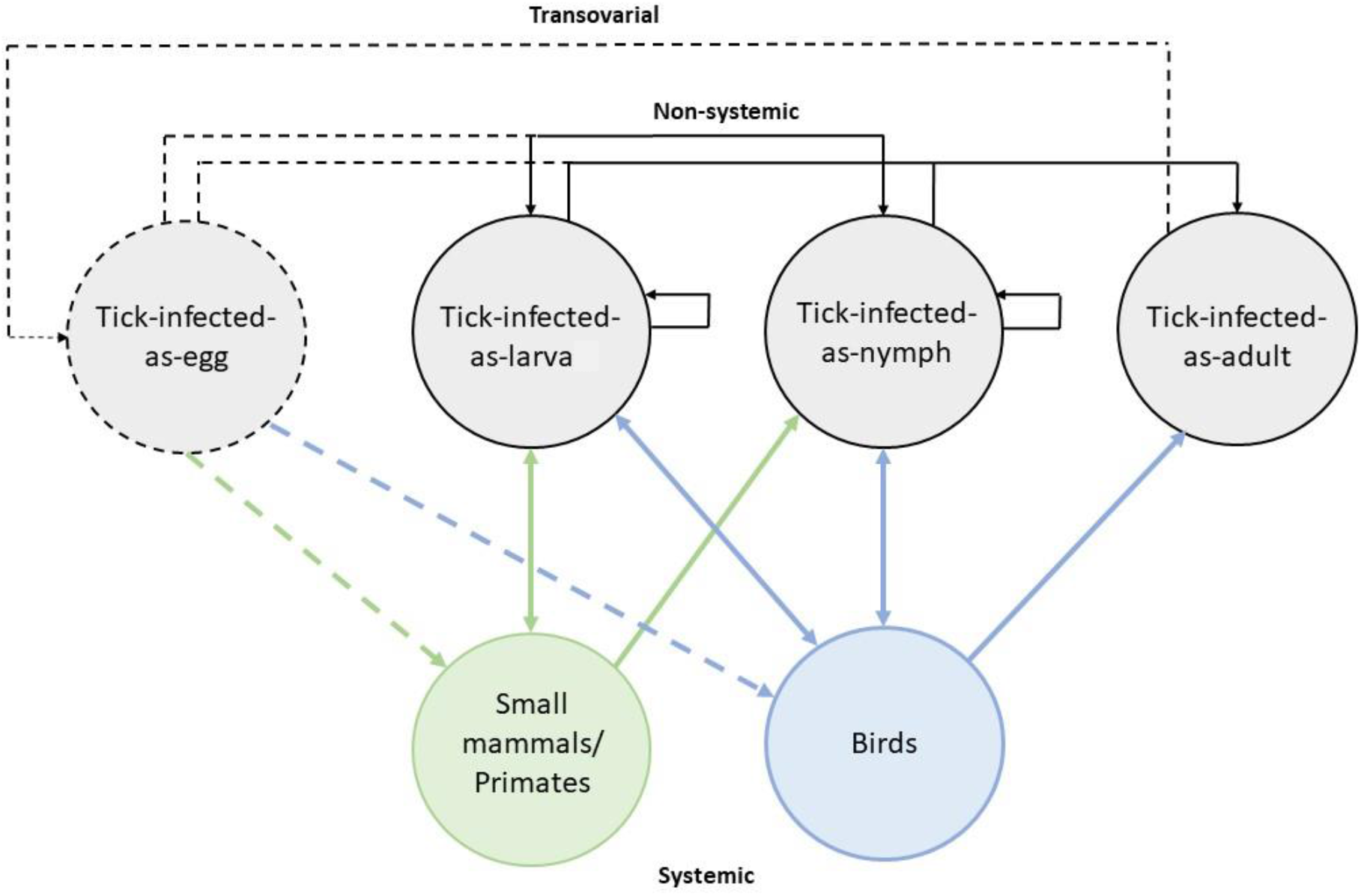
Transmission graph for both models. Solid lines show edges that are present in both models and dashed lines show edges that are only present when the tick-infected-as-egg type-at-infection and transovarial transmission are included in the model.

To adapt the model framework to explicitly consider different vertebrate host types we expand on previous equations outlined in Hartemink et al. (2008). In that previous model, the *H_c_* parameter acts as a proxy for the probability that a tick will bite a host that is competent for transmission. In this model, we include an additional parameter *Pb_i_* as a proxy for the probability that a tick bite is on competent host *i*, which is proportional to the proportion of the competent host community that consists of host *i*. This parameter was also made life stage specific to account for the assumption that adult ticks do not feed on small mammals or primates (Rajagopalan et al., 1968; Trapido et al., 1964). For example, the number of birds infected by a tick that had been infected as a nymph would be described by Eq.1.

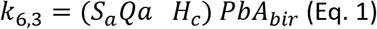

Where, *S_α_* is the probability of a nymph surviving to become an adult and biting any host. *Qa* is the probability of transmission, given that an adult tick bites a host and *PbA_bir_* is the probability that an adult tick will bite a bird, that is based on the relative host density of birds and the preference of the relevant tick life stage to bite a bird (see also Supplementary information S1.1)

By explicitly considering each host, we are then able to calculate the relative contribution of each host to systemic transmission. The relative contribution of systemic transmission to *R*_0_ can be calculated by composite elasticities of matrix elements associated with all systemically infected hosts, as outlined in Hartemink et al. (2008) and Matser et al. (2009). In this framework, each host is associated with its own matrix elements allowing the relative contribution of each vertebrate host to systemic transmission to be established by calculating the composite elasticities of matrix elements associated with each individual host.

### Scaling tick abundance and burden

We aim for our models to reflect the fact that tick densities may vary between areas, e.g. as a result of different habitat or different availability of hosts. Tick densities may affect tick burdens on hosts, as well as the number of eggs produced by female ticks. To account for factors influencing tick abundance in the environment and tick burden on hosts, we used two approaches, namely two different scaling factors to reduce or increase the number of eggs produced by adult females and tick burdens on hosts, and we compared the results. These scaling factors influence the number of tick-infected-as-egg produced as a result of transovarial transmission, as well as the number of each life stage present and co-feeding on hosts.

The first scaling factor was used to account for the potential influence of cattle density on tick abundance and burden. As there are no data available from India on the rate of increase in tick abundance as a result of increased cattle density, we use results from a different disease system that outlines the rate of increase in the density of questing nymphs in relation to the density of deer (0.026) (Dickinson et al., 2020). This scaling factor was applied to the number of eggs produced by adult females and to all parameters related to tick burden (Eq.2).

The second scaling factor accounts for differences in tick densities in different habitat types. In this case we assume that tick abundance and burden on hosts will vary with habitat type as result of availability of hosts and environmental factors that may influence the survival of ticks. This scaling factor was calculated as one minus the proportion decrease in tick density in relation to the habitat type with the highest tick density. As above, it was applied to the number of eggs produced by adult females and to all parameters related to tick burden (Eq.3). To estimate the change in tick abundance across habitat types we used data on the difference between numbers of nymphs collected in different land use types in Europe (Rosà et al., 2018). This provides some estimate of how ticks may be distributed across habitat types in general but it is worth noting that agricultural practices and plants used in crops and plantations will differ and the focal area in India has a greater number of forest types being in a tropical region. The reason for using data from Europe was due the lack of availability of local data on the distribution of *Haemaphysalis* ticks across different land use types.

Cattle is not a type-at-infection in the matrix model, as it is a dead end host for KFD virus. It was however as part of the cofeeding matrix elements within the model, to account for the fact that cofeeding can take place on a cow. For example, the formula for the matrix element quantifying the number of adult ticks infected by a tick that was infected as a nymph through co-feeding is described by Eq.2 (using scaling factor calculated using cattle density) and Eq. 3 (using scaling factor calculated using habitat type):

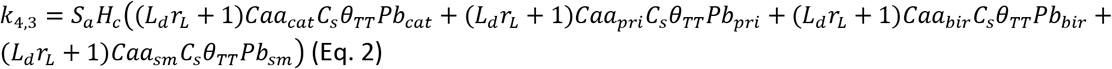

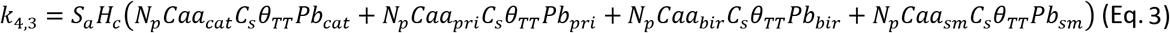

Where *Pb_sm_, Pb_pri_*, *Pb_bir_* and *Pb_cat_* outline the proportion of the host community competent for non-systemic transmission that is made up of small mammals, primates, birds and cattle respectively. *θ_TT_* is the efficiency of transmission from tick to tick and *Cαα_i_*, is the number of adult ticks expected to be feeding with a single adult tick on each host species. *L_d_* and *r_L_*(Eq.2) are the density of cattle and the rate of increase in tick burden respectively. *N_p_* is one minus the proportion of decrease in tick abundance in relation to the habitat type with the highest tick abundance. For all equations and parameters used in matrix see Supplementary Material S1.

### Scenarios

Using this NGM framework, we outlined different scenarios to see how R_0_ and the contribution of different routes of transmission and vertebrate host to R_0_ may change in different habitat types with different host community compositions. To do this we focused on five different relevant habitat types present in the agro-forestry mosaic in which KFD spillover occurs (Pattnaik, 2006; Purse et al., 2020): evergreen forest, deciduous forest, agriculture and plantations (excluding paddy), paddy and areas of human habitation. In each habitat type we varied parameters within the model to reflect potential habitat-specific differences in host community composition. Data on host community composition in each habitat type is not available at this time from specific KFD affected areas within the Western Ghats. However, we were able to source data from studies carried out in Western Ghats that report the number of animals recorded or trapped and the area covered (Bapureddy et al., 2015; Kumara & Singh, 2004; Molur & Singh, 2009; Pramod et al., 1997). We used this data as a proxy for small mammal, bird, primate and cattle densities in different habitat types in KFD affected areas.

For small mammals, Molur & Singh (2009) recorded the number of individuals trapped and the area trapped (ha) in evergreen forest, deciduous forest, agriculture (including ginger and paddy), plantations (banana, cardamom, coffee, orange and tea) and areas of human habitation. In this case, we scaled the number of individuals trapped per hectare up to the number of individuals trapped per 1 km^2^ in evergreen and deciduous forest, agriculture (including paddy) and human habitation habitat types. We also used this method for plantation habitat types: banana, cardamom, coffee, orange and tea and calculated the mean number of individuals trapped per 1 km^2^ to estimate small mammal density in plantations.

For birds, Pramod et al. (1999) outline data for number of birds in evergreen forest, deciduous forest, plantation, paddy and areas of human habitation in the Western Ghats. Abundance data for birds was collected in 600m x 100 m belts transects in different habitat types (Kunte et al., 1999; Pramod et al., 1997). We scaled this data up to individuals recorded per km^2^ assuming transects covered an area of 0.06 km^2^. As a proxy for bird density in evergreen forest, we used the middle of the range of densities of birds per km^2^ in both the evergreen forest and semi-evergreen forest habitat types. For deciduous forest, we used the moist and dry deciduous forests habitat types. Data was also available for individuals recorded in paddy fields, monoculture plantations habitat types as well as the garden and habitation habitat type. To refine our estimates of bird densities we focused on bird species that tend to live or forage on the ground and which tended to have the highest burden of ticks in KFD areas (Rajagopalan, 1972). This included babblers, thrush, mynas, crow pheasant and junglefowl (See Supplementary material S1.4 for list of focal species). We utilised bird detection frequency data from eBird (eBird, 2021) for Karnataka in India calculating the sum of the mean detection frequency for all species and the proportion of that made up by our focal bird species. We then scaled bird density estimates based on this proportion.

For primates, Bapureddy et al. (2015) report estimated density of individuals per km^2^ for evergreen and deciduous forest types at 12 locations. Using this data we calculated the mean density in locations where habitat types included deciduous forest type or evergreen forest. (M. Singh & Rao, 2004) report the number of individuals per km of occupied area in intensive agriculture (including paddies) and dry agriculture (including plantations) for three years (1989, 1998 and 2003) and we used the mean across the three time periods to inform density of primates in these habitat types.

As proxy for cattle densities we used the mean indigenous cattle density across the Shimoga region in Karnataka (60 cattle per km) (Purse et al., 2020) and adjusted this using recently collected data on habitat use of indigenous cattle in Shimoga (Kumar et al. unpublished). This data on cattle movement outlines the proportion of time spent in a habitat type vs the proportion of area made up of that habitat type, 0.67 (evergreen forest), 1.99 (deciduous forest), 1.14 (plantation) and 0.74 (cropland). Therefore, we scaled density to reflect the ratio of the proportion of time spent in each habitat type vs the proportion of habitat available in the area and reduced density where the ratio was less than one: 40.2 cattle per km (evergreen forest), 60 cattle per km (deciduous forest), 60 cattle per km (plantation) and 44.4 cattle per km (cropland).

The densities calculated above were used as a proxy for the number of hosts per 1 km^2^ and proportion of the total host community that each host makes up (Table 1). (See supplementary material S2 for parameters for each scenario). We performed 100 runs of the model for each scenario, using either cattle density or habitat type as a scaling factor.

**Table 1:** Table showing proportion of host community made up of small mammals (sm), birds (bir), primates (pri) and cattle (cat) in each habitat type. Numbers in brackets show estimated density (individuals per km^2^).

| Parameter | Evergreen forests | Deciduous forests | Agriculture /plantation | Paddy | Human habitation |
| --- | --- | --- | --- | --- | --- |
| $Pb_{sm}$ | 0.922<br>(1537.65) | 0.947<br>(2891.56) | 0.913<br>(1283.42) | 0.933<br>(1760) | 0.92<br>(2315.78) |
| $Pb_{bir}$ | 0.045<br>(74.93) | 0.030<br>(93.06) | 0.041<br>(57.53) | 0.04<br>(75.24) | 0.046<br>(116.6) |
| $Pb_{pri}$ | 0.009<br>(15.25) | 0.008<br>(25.6) | 0.004<br>(5.23) | 0.003<br>(5.86) | 0.010<br>(25.6) |
| $Pb_{cat}$ | 0.024<br>(40.1) | 0.020<br>(60) | 0.043<br>(60) | 0.024<br>(44.4) | 0.024<br>(60) |

### Sensitivity and elasticity analysis

For each run, the sensitivity of *R*_0_ to changes in individual parameters (*S*) was calculated using the equation defined in (Caswell, 2001).

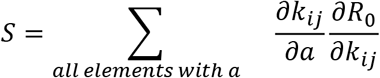

With 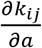 representing the change of the focal element (*k_ij_*) with respect to the focal parameter *a* multiplied by the sensitivity of *R*_0_ to the focal element (*k_ij_*). The sum of these values for all matrix elements derived using the focal parameter (*a*) gives the sensitivity of *R*_0_ to each individual parameter. The elasticity can then be calculated by 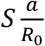.

## Results

We performed 100 runs for each scenario with the model using cattle density as a scaling factor (“cattle model”) and the model using habitat type as a scaling factor (“habitat model”), both including and excluding transovarial transmission. The results from these models show that in all scenarios the models excluding transovarial transmission had an *R*_0_ of less than one in the majority of runs when habitat type (Mean R_0_= 0.071, SD=0.035) and cattle density (Mean R_0_=0.135, SD=0.06) were used as scaling factors (Figure 2 & Figure S2.1). When transovarial transmission is included in the model using cattle density to scale tick abundance and burden *R*_0_ was similar and consistently exceeded one in all scenarios (MeanR0 =1.83, SD=0.789) (Figure S2.1). However, when tick abundance and burden is scaled based on the habitat type, there was more variation in *R*_0_, which was generally below one in the human habitation, paddy and plantation and above one in evergreen forest and deciduous forest (Mean R_0_= 1.05, SD=0.49) (Figure 2).

**Figure 2:**
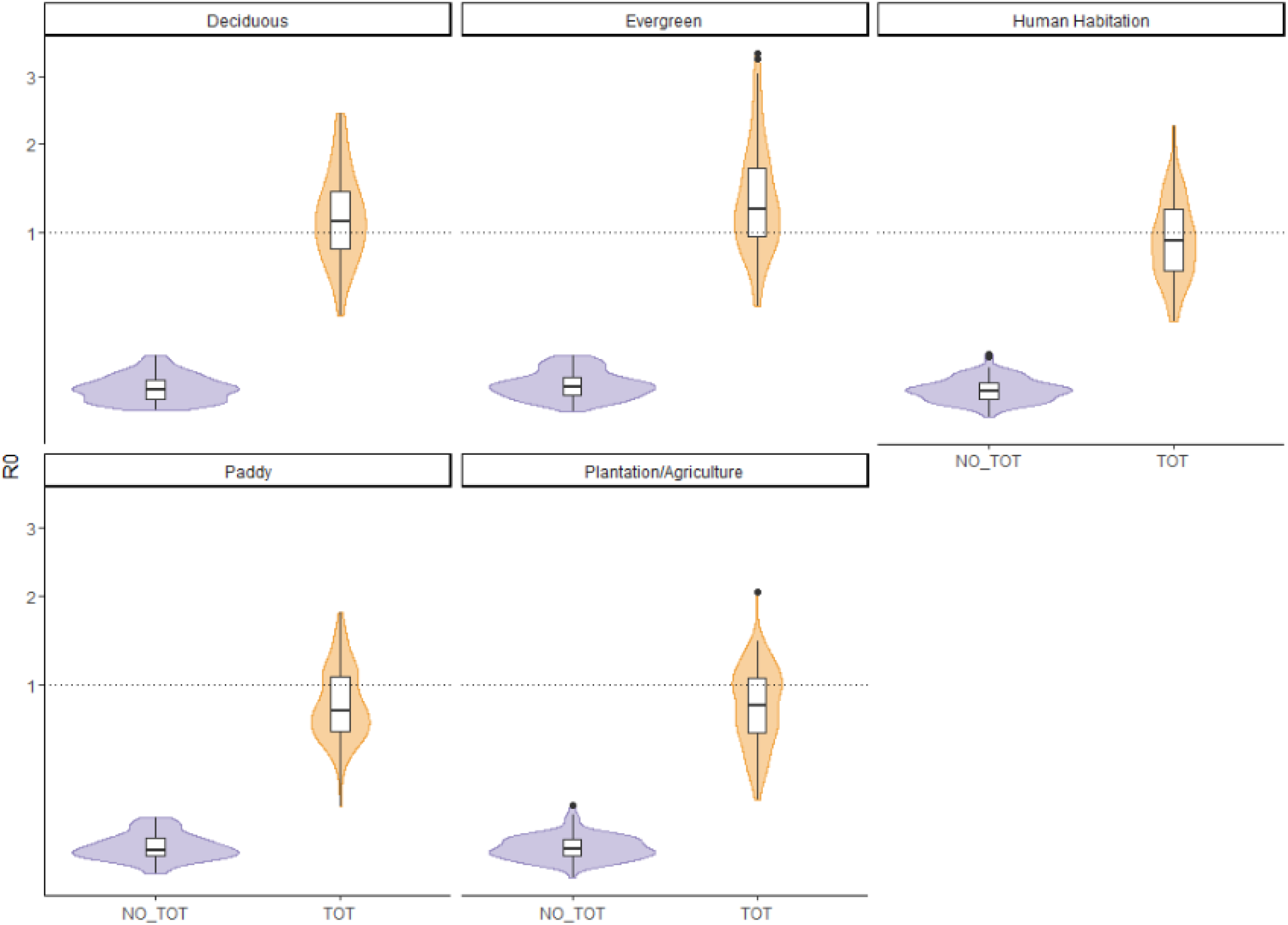
Distribution of values in each habitat type for models using scaling factor based on habitat type. Plot shows results for models excluding transovarial transmission (NO_TOT) and including transovarial transmission (TOT). For results using cattle density as scaling factor see Supplementary material S2 Figure S2.1

Across all models and scenarios, we found that the relative contributions to R_0_ for systemic transmission (Mean composite elasticity=0.48, SD=0.11) and transovarial transmission (0.46, SD=0.08) were quite similar, and much higher than for non-systemic transmission (0.04, SD=0.04) (Figure 3 & Figure S2.2). In the model without transovarial transmission, almost all transmission was systemic (Mean composite elasticity=0.965, SD=0.04), with a very small contribution to R_0_ from non-systemic transmission (Mean composite elasticity=0.03, SD=0.04) (Figure S2.4).

**Figure 3:**
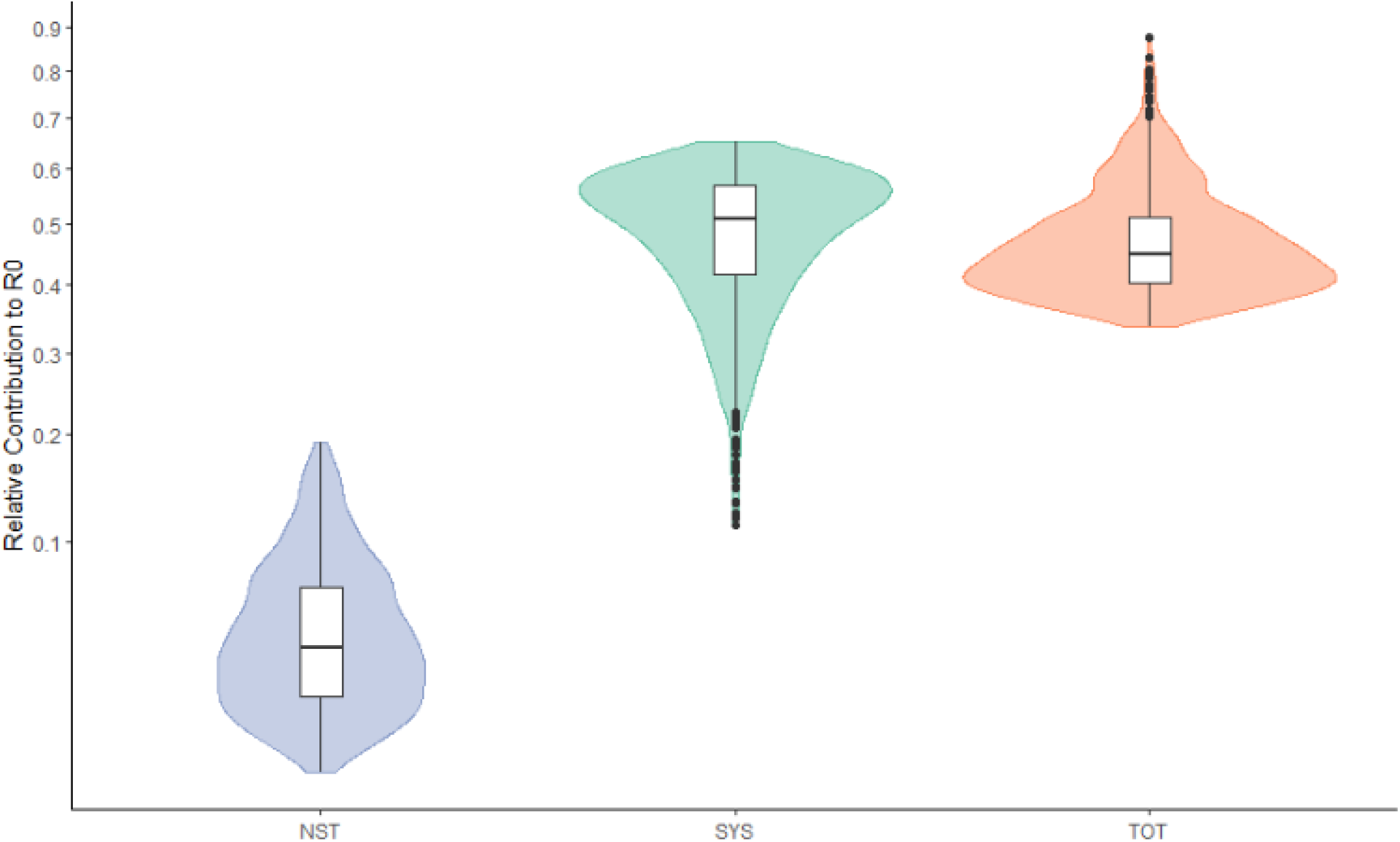
Distribution of values for relative contribution to R_0_ for each route of transmission across all scenarios and different scaling factors with transovarial transmission included in the model. (NST= Non-systemic transmission, SYS= Systemic transmission,TOT=transovarial). For results broken down by habitat type and scaling factor see Supplementary material S2 Figure S2.2.

By further breaking down systemic transmission it is clear that there is a marked difference between the relative contribution of small mammals, birds and primates to systemic transmission in both models (Figure 4). Small mammals consistently had the greatest contribution to systemic transmission (Mean relative contribution to systemic transmission= 0.85, SD=0.12) followed by birds (Mean relative contribution to systemic transmission= 0.12, SD=0.11) and finally primates (Mean relative contribution to systemic transmission= 0.02, 0.02).

**Figure 4:**
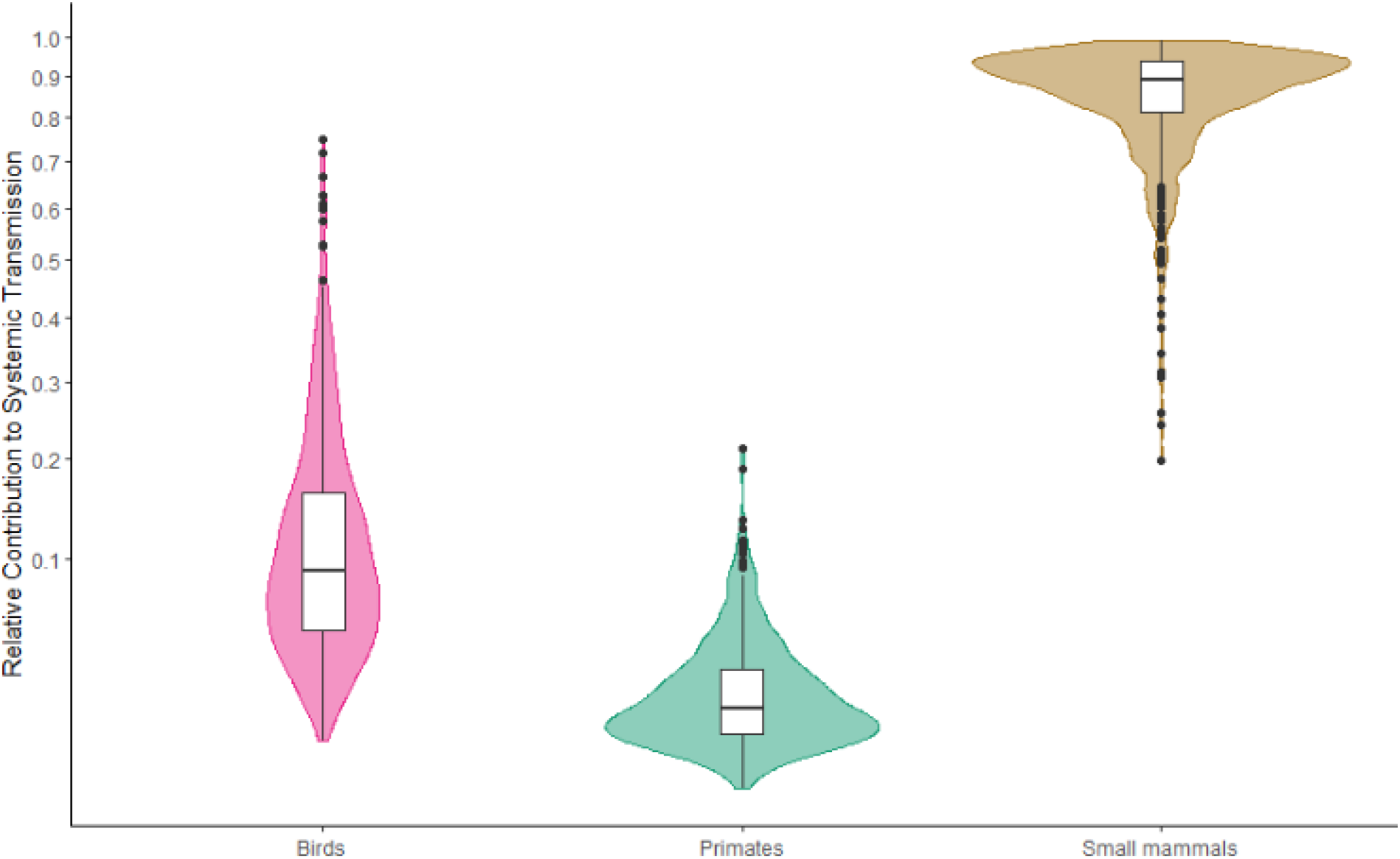
Distribution of values for relative contribution to systemic transmission for each host across all scenarios and different scaling factors with transovarial transmission included in the model. For results broken down by habitat type and scaling factor see Supplementary material S2 Figure S2.3.

Extracting the elasticity of individual matrix elements revealed that the number of eggs infected by tick-infected-as-nymph and the number of eggs infected by tick-infected-as-egg (i.e. from their mother) were among the matrix elements that had the highest relative contribution to R_0_. The number of small mammals infected by ticks-infected-as-eggs and the number of nymphs infected by small mammals also had a high relative contribution to R_0_. (Table 2).

**Table 2:** Table shows the relative contribution to R_0_ from individual matrix elements (mean with 2.5% and 97.5% quantiles) and description of the contribution of matrix elements to number of infecteds for both the model using cattle density as a scaling factor and habitat type as a scaling factor. Matrix elements with a contribution of above 0.01 are included.

| **Matrix index** | **Description** | **Cattle Density** | **Habitat** |
| --- | --- | --- | --- |
| k13 | Number of eggs infected by tick-infected-as-nymph | 0.23 (0.18-0.29) | 0.22 (0.17-0.27) |
| k51 | Number of small mammal infected by tick-infected-as-egg | 0.2 (0.15-0.25) | 0.21 (0.16-0.27) |
| k11 | Number of eggs infected by tick-infected-as-egg | 0.17 (0.08-0.26) | 0.19 (0.09-0.27) |
| k35 | Number of nymphs infected by an infected small mammal | 0.17 (0.11-0.23) | 0.18 (0.12-0.24) |
| k12 | Number of eggs infected by tick-infected-as-larvae | 0.05 (0.03-0.08) | 0.05 (0.02-0.08) |
| k31 | Number of nymphs infected by tick-infected-as-egg | 0.05 (0.01-0.08) | 0.03 (0.01-0.04) |
| k25 | Number of larvae infected by an infected small mammal | 0.03 (0.01-0.06) | 0.03 (0.01-0.05) |
| k61 | Number of birds infected by tick-infected-as-egg | 0.03 (0.01-0.05) | 0.03 (0.01-0.04) |
| k26 | Number of larvae infected by an infected by bird | 0.02 (0-0.03) | 0.01 (0-0.02) |
| k14 | Number of eggs infected by a tick-infected-as-adult | 0.01 (0-0.02) | - |
| k36 | Number of nymphs infected by an infected bird | 0.01 (0-0.02) | 0.01 (0-0.02) |

The sensitivity of R_0_ to individual parameters was assessed for both models including transovarial transmission. This analysis demonstrated both high sensitivity and elasticity values for parameters within the models indicating that changes in some parameters can result in large changes to R_0_. In both models the rate of transovarial transmission (*R_a_*) had both high sensitivity and elasticity values. For the model with cattle density as a scaling factor, this analysis revealed relatively high sensitivity and elasticity values for *r_L_* (the rate of increase in tick abundance and burden with cattle density), *S_l_* (survival probability from egg to feeding larva), *S_n_* (survival probability from feeding larva to feeding nymph) and *S_a_* (survival probability from feeding nymph to feeding adult). These parameters also had relatively higher sensitivity values in the model using habitat type as a scaling factor with *S_l_*, S_n_and *S_a_* having relatively high values for sensitivity and elasticity values followed by *N_p_* (1 – the proportion decrease in tick abundance).

**Figure 4:**
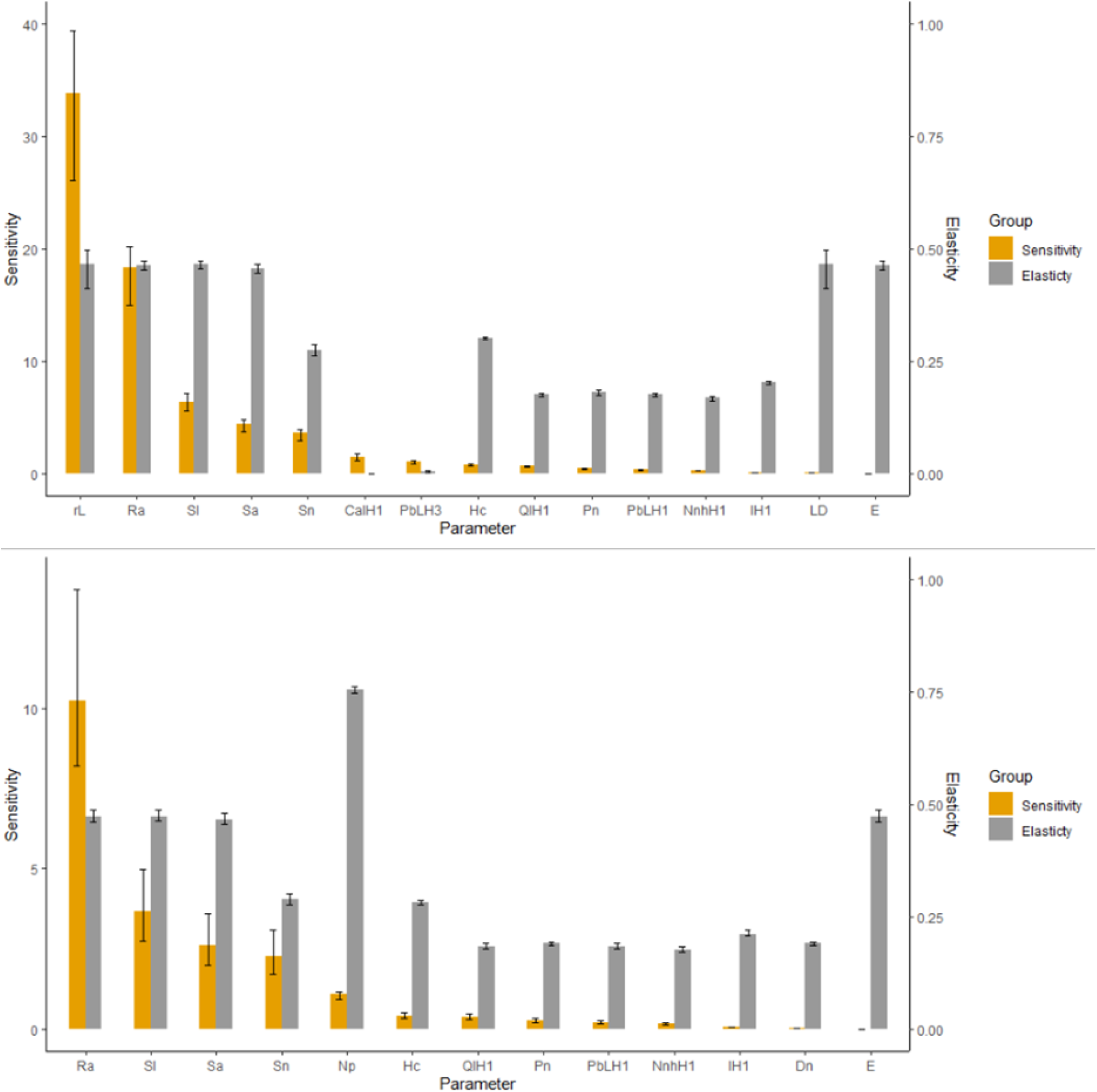
Mean sensitivity and elasticity values for individual parameters across all scenarios when transovarial transmission is included in the model. (top) model with cattle density as scaling factor and (bottom) model with habitat type as scaling factor. Error bars show 2.5% and 97.5% quantiles. Parameters were plotted if they had a sensitivity of greater that 1 or an elasticity of greater than 0.1.

## Discussion

This study developed a novel next generation matrix-modelling framework with wide applicability for improving understanding of the relative role of different hosts and transmission processes in the establishment of multi-host tick-borne pathogens in diverse and changing landscapes. When applied to the case study disease, Kyasanur Forest Disease, in south India, despite relatively sparse empirical data, this approach highlights that key transmission processes (transovarial transmission) and reservoir hosts (birds and small mammals) likely play the most important role in maintaining transmission across all habitat types regardless of the scaling factor used. This study also suggests potential variation in disease transmission risk between habitat types in the mosaic when habitat variation in tick abundance, tick burden and host density are accounted for. We thus recommend that the focus of research and management is extended beyond primates to the other competent hosts that may play a more important role in maintaining KFD and influencing the distribution and density of infected ticks. Here we set our findings in the context of prior empirical and modelling work for this and other tick-borne disease systems and identify key research priorities to inform interventions to reduce the burden of KFD in India.

Based on our findings and those of Burthe et al. (2021) it is becoming increasingly clear that there is a need to better evaluate the host preferences of *Haemaphysalis* ticks, the competence and susceptibility of hosts and subsequently the role of different hosts in influencing the maintenance of KFD and distribution of ticks. Understanding this at the local scale in areas prone to KFD outbreaks is also important as host range reported over larger regions may not necessarily translate to how ticks utilise hosts in a local context (McCoy et al., 2013). Much of the research on KFD and subsequent management recommendations have focused on the role of primates in creating hotspots for the transmission of KFD, with a focus on burning monkey carcasses and dusting insecticides within 50 feet of reported monkey deaths (Burthe et al., 2021). However, when accounting for both host density and tick burden on different hosts, our models suggest that small mammals and birds are likely to have the largest contribution to systemic transmission of KFD. Small mammals are an ideal reservoir for tick borne pathogens as an easily infected abundant host, which may not suffer from severe symptoms and regularly produces naive individuals (Michelitsch et al., 2019). Indeed, small mammals have already been highlighted as important reservoirs for several other tick borne pathogens with complex, multi-host transmission cycles including Lyme disease (Ostfeld, 1997), Rickettsial pathogens (Azad & Beard, 1998) and other tick-borne flaviviruses such as tick-borne encephalitis (Mlera & Bloom, 2018). Ground frequenting birds may also play an important role as an additional abundant and highly mobile host that could also introduce infected ticks to new areas (Brinkerhoff et al., 2019; Hasle, 2013; Loss et al., 2016; Ogden et al., 2008) and a phylogeographic analysis has already highlighted the potential role of migratory birds in carrying KFD infected ticks between Saudi Arabia and India (Brinkerhoff et al., 2019; Hasle, 2013; Mehla et al., 2009). Currently there is a lack of more recent data on the tick burdens on hosts and much of the available data on tick infestation of small mammals, birds and primates is from research carried out before 1972, after which further human modification of landscapes and habitats has taken place. Therefore, we need more up to date data on the distribution of potential hosts and on the tick burdens and the prevalence of KFD in small mammals, birds and primates to understand the contribution of these different hosts to maintenance of KFD and the distribution of ticks.

Additionally, we need a better understanding of the evolutionary relationships between KFD strains infecting these hosts versus those infecting humans in an effort to narrow down links between KFD strains infecting humans and strains circulating in wildlife reservoirs. Phylogenetic studies have regularly revealed that pathogen species once thought to be generalists can exhibit cryptic host specificity, with individual strains exhibiting strong host associations, either as a result of compatibility barriers or encounter barriers (Withenshaw et al., 2016). For example, evidence of limited cross-species transmission of rabies lineages in bats and strong host associations of *Borrelia burgdorferi* strains circulating in bank voles and chipmunks in France (Jacquot et al., 2014; Streicker et al., 2010). Evidence from a study investigating strains infecting primates, humans, ticks and rodents from the genus *Rattus* has so far demonstrated little divergence between strains infecting these different hosts, indicating a lack of host associations, but further studies would help to shed light on links between strains circulating in other wildlife reservoirs and strains infecting humans. Another recent study has also provided insights into evolutionary relationships between KFD strains found in ticks, primates and humans and the likely pattern of spread of KFD across Karnataka since 1957(Yadav et al., 2020). This study attributed the distribution of KFD across Karnataka to the movement of tick-infested primates moving through forests, however, as of yet there is no information on the evolutionary relationships between human and primate strains and strains infecting small mammals, birds and ticks on cattle. Once again, further investigation is needed to understand the spread of KFD in India since it is just as plausible that longer distance dispersal of ticks infected with KFD could occur through movements of birds or cattle (Brinkerhoff et al., 2019; Hasle, 2013; Mehla et al., 2009). The later of of which could feasibly be subject to interventions aimed at limiting spread to new areas (e.g. use of repellents and pre- and post-movement tick removal on cattle) (Burthe et al., 2021)

At the moment there is no evidence that cattle are systemically infected with KFD but they are parasitised by *Haemaphysalis* ticks and may provide opportunities for transmission between ticks through non-systemic transmission or the movement or maintenance of infected adult tick populations (Balasubramanian et al., 2019). Consistent with previous work and unusually among well-studied tick-borne pathogens (Matser et al., 2009), our models suggest that transovarial transmission plays an important role in the maintenance of KFD with a marked decrease in R_0_ (below 1) when transovarial transmission is excluded. Despite being demonstrated in laboratory studies using *Ixodes petauristae* (Singh et al., 1962, 1968) there is still a lack of evidence that transovarial transmission of KFD occurs in the wild in *Haemaphysalis* ticks. However, the models used in this study indicate the potential for ticks to act as a reservoir for KFD and highlight the importance of small mammals being infected by larvae infected via transovarial transmission, when transovarial transmission occurs. This is an important question for understanding the maintenance of KFD as our sensitivity analysis highlights that R_0_ has a high sensitivity to the rate of transovarial transmission and also survival of different tick life stages. If ticks do play a significant role in maintaining KFD, cattle may play an important role by acting as widely available reproductive hosts for female adult ticks and influencing abundance and distribution of ticks. A number of other studies have highlighted similar roles of larger mammals in tick-borne disease systems in Europe, showing the important role of deer influencing abundance and distribution of *I. ricinus* in Europe (Dickinson et al., 2020; Gilbert et al., 2012; Hofmeester et al., 2017) and the potential for non-systemic transmission through co-feeding on sheep (a non-competent host) in maintaining *B. burgdorferi (Ogden et al., 1997).* The limited empirical data on the number and life stage ticks parasitising cattle in KFD-affected regions has also been largely collected in agricultural settings (Balasubramanian et al., 2019) rather than in the specific agro-forest mosaics in which spillover is occurring. When looking at the sensitivity of R_0_ to the rate of increase in tick abundance due to cattle density it is clear that the model including cattle density as a scaling factor had high sensitivity to this parameter. This further highlights the need to determine how cattle density and movement influences abundance and distribution of ticks to which humans are exposed to inform future models, which would be facilitated by sampling of ticks across different habitat types at the human-animal-forest interface and linking this to frequency of use of these habitat types by cattle and humans.

It is also apparent that our model had relatively high sensitivity to the habitat scaling factor compared to other parameters included in the model. It is well established that the survival, abundance and distribution of ticks is dependent on the overlap of suitable off-host environmental conditions and the distributions and movements of hosts (Diuk-Wasser et al., 2021; Leal et al., 2020; Pfäffle et al., 2013). There have been many studies in Europe that have found that habitat type or composition can influence the abundance of different tick species and prevalence of associated pathogens such as *B. burgdorferi* and *Anaplasma phagocytophilum* (Ehrmann et al., 2017, 2018; Rosà et al., 2018; Tack et al., 2012). Our models did show some variation in R_0_ across different land use classes when we assume that tick abundance, burden on hosts and host density varies with habitat type. These findings suggest that forested areas are more likely to have higher numbers of infected ticks and vertebrate hosts compared to areas of agriculture and plantations, paddy and human habitation. This is consistent with previous findings that visits to forests, grazing of cattle in forests and handling of dry leaves are significant risk factors for contracting KFD in India (Kasabi et al., 2013). However, it should be noted that our model is currently informed by the relative abundance of *Ixodes ricinus* ticks in different land use types in Europe (Rosà et al., 2018) because we still have a limited understanding of how the abundance of *Haemaphysalis* ticks in KFD-affected areas varies across habitat types. It is understood that this genus of ticks generally prefers forested habitats (Geevarghese & Mishra, 2011) but our understanding of how human modifications to landscapes have altered suitability for ticks across landscapes is still limited and tick bite exposure may not necessarily be limited to the primary habitats of ticks (Vanwambeke & Schimit, 2021). A better understanding of risk in different habitat types can be gained by gathering empirical data on the abundance of questing ticks in these habitats and the prevalence of KFD in ticks. In particular, previous studies have highlighted evidence that KFD may be an ecotonal disease arising as a result of human encroachment into forested areas (Purse et al., 2020). Modelling of historical outbreak patterns (2014-2020) indicates that spillover events to humans were more likely in diverse forest mosaics with high proportions of moist evergreen and plantation habitats (Purse et al., 2020). Therefore, sampling tick populations in different habitats would ideally be accompanied by sampling across ecotones, for example from forests across ecotones and into matrix habitat (Despommier et al., 2007; Diuk-Wasser et al., 2021), to improve our understanding of how the abundance and distribution of ticks varies across agro-forest mosaics. Ideally, this data should also be combined with data on human use of landscapes and coping capacity to shed light on how tick bite risk varies across focal areas (Vanwambeke & Schimit, 2021).

Non-systemic transmission through co-feeding has been highlighted as playing an important role in the maintenance of flaviviruses such as tick-borne encephalitis virus in Europe (Matser et al., 2009; S E Randolph et al., 1996). In contrast, our model currently suggests that this form of transmission has a relatively low contribution to R_0_ for KFD, consistent with previous modelling of this system by Matser et al., (2009). The efficiency of non-systemic transmission can be dependent on the distance between feeding ticks, with efficiency shown to drop at distances of greater than 1cm (between co-feeding individuals) for *B. burgdorferi (Richter et al., 2002)* and the common aggregation of ticks within distances of around 1cm being thought to be important for transmission of tick borne-encephalitis (Randolph, 2011). Seasonality of activity of immature tick life stages can also influence non-systemic transmission between life stages, with seasonal synchrony of the activity of larvae and nymphs thought to be important in the transmission of TBEV in Europe (Randolph et al., 1999). In Karnataka, previous data from blanket drags and the prevalence of ticks on small mammals suggests that larval ticks are most abundant in November and December whilst nymphs are most abundant in January and February (Rajagopalan et al., 1968), highlighting potential for lack of synchrony for immature life stages and estimates for aggregation of ticks on hosts was not available for this system at this time. However, collection of up to date data on seasonality of different tick species, and their frequency and aggregation of ticks on hosts is ongoing, which will allow a better understanding of the potential contribution of non-systemic transmission to the maintenance of KFD.

This new framework can be readily applied to other tick-borne disease systems to explore the relative role of different types of transmission and hosts and allows the incorporation of variation in tick abundance and burden that may arise through habitat suitability and changing densities of important blood meal hosts. It can also aid in prioritising future research to reduce uncertainty about the processes and mechanisms involved in the maintenance of tick-borne pathogens. Using this novel next generation modelling approach, we have highlighted a number of critical knowledge gaps that, if filled, would greatly improve our understanding of the ecology and maintenance of KFD in the context of agro-forest mosaics. There is a great need to improve our knowledge of the spatio-temporal distribution of ticks and hosts within these modified landscapes and to better understand how ticks are likely to utilise these hosts. Coupling this with information on prevalence of KFD and understanding the evolutionary relationships between any KFD variants will be crucial to developing improved mechanistic modelling approaches that can be used to link human use of landscapes with natural foci of this pathogen for informing effective intervention strategies.

## Acknowledgements

We greatly appreciate the data on cattle habitat use in KFD-affected areas that was shared by the Ashoka Trust for Research in Ecology and the Environment, India (ATREE) and the Monkey Fever Risk project. We also appreciate the insightful discussions with researchers from ATREE and other members of the MonkeyFeverRisk and IndiaZooRisk research teams that helped to inform the development of these models.

