## Supplementary Information S1 for "Using mechanistic models to highlight research priorities for tick-borne zoonotic diseases: Improving our understanding of the ecology and maintenance of Kyasanur Forest Disease in India"

**Supplementary material S1**

**S1.1 Parameters used in next generation matrix models:**

**Parameters for models (excluding scaling factors)**

Parameters used for each scenario. Numbers separated by “;” indicate low and high end of the range.

Cofeeding parameters are derived using frequency data available for birds, primates and preliminary frequency data on the number of ticks found on rodents in Shimoga in India (Monkey Fever Risk Project). We generated distributions using joint probabilities and used the upper 95% quantile to define an upper limit for cofeeding parameters, with the lower limit of all cofeeding parameters being set 0.01. For example, the upper limit for the range of mean number of nymphs cofeeding with a single larva was derived using a distribution of probabilities calculated by multiplying the probability of having a single larva and the probabilities of having any number of nymphs. This range is then scaled using the Cs (fraction of ticks feeding close enough for non-systemic transmission).

Cofeeding numbers for cattle were informed by numbers of ticks of each life stage divided by number of cattle infested with ticks (168) from Balasubramanian (2019), which suggest only nymphs and adults were found on cattle. The average number of ticks feeding on individuals is 0.4 for nymphs and 1.87 for adults. As frequency data was not available for cattle we used the average number of ticks of each life stage per individual as the upper limit for cofeeding, which was then scaled using the Cs parameters.

| Parameter | Description | Evergreen forest | Deciduous forest | Agriculture/ plantation | Paddy | Human habitation | Reference |
|---|---|---|---|---|---|---|---|
| PbH1 | Proportion of host community made up of small mammals | 0.922 ; 0.922 | 0.942 ; 0.942 | 0.913 ; 0.913 | 0.933 ; 0.933 | 0.92 ; 0.92 | 1 |
| PbH2 | Proportion of host community made up of birds | 0.045 ; 0.045 | 0.03 ; 0.03 | 0.041 ; 0.041 | 0.04 ; 0.04 | 0.046 ; 0.046 | 2,3 |
| PbH3 | Proportion of host community made up of primates | 0.009 ; 0.009 | 0.008 ; 0.008 | 0.004 ; 0.004 | 0.003 ; 0.003 | 0.01 ; 0.01 | 4,5 |
| PbH4 | Proportion of host community made up of cattle | 0.024 ; 0.024 | 0.02 ; 0.02 | 0.043 ; 0.043 | 0.024 ; 0.024 | 0.024 ; 0.024 | 6 |
| PbLH1 | Proportion of host community made up of small mammals (larvae) | 0.922 ; 0.922 | 0.942 ; 0.942 | 0.913 ; 0.913 | 0.933 ; 0.933 | 0.92 ; 0.92 | 1 |

| | | | | | | | |
|---|---|---|---|---|---|---|---|
| PbLH2 | Proportion of host community made up of birds(larvae) | 0.045 ; 0.045 | 0.03 ; 0.03 | 0.041 ; 0.041 | 0.04 ; 0.04 | 0.046 ; 0.046 | 2,3 |
| PbLH3 | Proportion of host community made up of primates(larvae) | 0.009 ; 0.009 | 0.008 ; 0.008 | 0.004 ; 0.004 | 0.003 ; 0.003 | 0.01 ; 0.01 | 4,5 |
| PbLH4 | Proportion of host community made up of cattle(larvae) | 0.024 ; 0.024 | 0.02 ; 0.02 | 0.043 ; 0.043 | 0.024 ; 0.024 | 0.024 ; 0.024 | 6 |
| PbNH1 | Proportion of host community made up of small mammals(nymphs) | 0.922 ; 0.922 | 0.942 ; 0.942 | 0.913 ; 0.913 | 0.933 ; 0.933 | 0.92 ; 0.92 | 1 |
| PbNH2 | Proportion of host community made up of birds(nymphs) | 0.045 ; 0.045 | 0.03 ; 0.03 | 0.041 ; 0.041 | 0.04 ; 0.04 | 0.046 ; 0.046 | 2,3 |
| PbNH3 | Proportion of host community made up of primates(nymphs) | 0.009 ; 0.009 | 0.008 ; 0.008 | 0.004 ; 0.004 | 0.003 ; 0.003 | 0.01 ; 0.01 | 4,5 |
| PbNH4 | Proportion of host community made up of cattle(nymphs) | 0.024 ; 0.024 | 0.02 ; 0.02 | 0.043 ; 0.043 | 0.024 ; 0.024 | 0.024 ; 0.024 | 6 |
| PbAH1 | Proportion of host community made up of small mammals (adults) | 0 ; 0 | 0 ; 0 | 0 ; 0 | 0 ; 0 | 0 ; 0 | 1 |
| PbAH2 | Proportion of host community made up of birds (adults) | 0.045 ; 0.045 | 0.03 ; 0.03 | 0.041 ; 0.041 | 0.04 ; 0.04 | 0.046 ; 0.046 | 2,3 |
| PbAH3 | Proportion of host community made up of primates (adults) | 0 ; 0 | 0 ; 0 | 0 ; 0 | 0 ; 0 | 0 ; 0 | 4,5 |
| PbAH4 | Proportion of host community made up of cattle (adults) | 0.024 ; 0.024 | 0.02 ; 0.02 | 0.043 ; 0.043 | 0.024 ; 0.024 | 0.024 ; 0.024 | 6 |
| Sl | Survival probabilities from egg to feeding larva | 0.05 ; 0.25 | 0.05 ; 0.25 | 0.05 ; 0.25 | 0.05 ; 0.25 | 0.05 ; 0.25 | 7 |
| Sn | Survival probabilities from larva to feeding nymph | 0.05 ; 0.25 | 0.05 ; 0.25 | 0.05 ; 0.25 | 0.05 ; 0.25 | 0.05 ; 0.25 | 7 |
| Sa | Survival probabilities from nymph to feeding adult | 0.15 ; 0.25 | 0.15 ; 0.25 | 0.15 ; 0.25 | 0.15 ; 0.25 | 0.15 ; 0.25 | 7 |
| CllH1 | # of larvae co-feeding with a larva (small mammals) | 0.01 ; 0.49 | 0.01 ; 0.49 | 0.01 ; 0.49 | 0.01 ; 0.49 | 0.01 ; 0.49 | 8 |
| CnlH1 | # of nymphs co-feeding with a larva (small mammals) | 0.01 ; 0.52 | 0.01 ; 0.52 | 0.01 ; 0.52 | 0.01 ; 0.52 | 0.01 ; 0.52 | 8 |

| | | | | | | | |
|---|---|---|---|---|---|---|---|
| CalH1 | # of adults co-feeding with a larva (small mammals) | 0 | 0 | 0 | 0 | 0 | 8 |
| ClnH1 | # of larvae co-feeding with a nymph (small mammals) | 0.01 ; 0.49 | 0.01 ; 0.49 | 0.01 ; 0.49 | 0.01 ; 0.49 | 0.01 ; 0.49 | 8 |
| CnnH1 | # of nymphs co-feeding with a nymph (small mammals) | 0.01 ; 0.55 | 0.01 ; 0.55 | 0.01 ; 0.55 | 0.01 ; 0.55 | 0.01 ; 0.55 | 8 |
| CanH1 | # of adults co-feeding with a nymph (small mammals) | 0 | 0 | 0 | 0 | 0 | 8 |
| ClaH1 | # of larvae co-feeding with an adult (small mammals) | 0 | 0 | 0 | 0 | 0 | 8 |
| CnaH1 | # of nymphs co-feeding with an adult (small mammals) | 0 | 0 | 0 | 0 | 0 | 8 |
| CaaH1 | # of adults co-feeding with an adult (small mammals) | 0 | 0 | 0 | 0 | 0 | 8 |
| CllH2 | # of larvae co-feeding with a larva (birds) | 0.01 ; 3.26 | 0.01 ; 3.26 | 0.01 ; 3.26 | 0.01 ; 3.26 | 0.01 ; 3.26 | 9 |
| CnlH2 | # of nymphs co-feeding with a larva (birds) | 0.01 ; 0.12 | 0.01 ; 0.12 | 0.01 ; 0.12 | 0.01 ; 0.12 | 0.01 ; 0.12 | 9 |
| CalH2 | # of adults co-feeding with a larva (birds) | 0.01 ; 0.039 | 0.01 ; 0.039 | 0.01 ; 0.039 | 0.01 ; 0.039 | 0.01 ; 0.039 | 9 |
| ClnH2 | # of larvae co-feeding with a nymph (birds) | 0.01 ; 0.71 | 0.01 ; 0.71 | 0.01 ; 0.71 | 0.01 ; 0.71 | 0.01 ; 0.71 | 9 |
| CnnH2 | # of nymphs co-feeding with a nymph (birds) | 0.01 ; 0.16 | 0.01 ; 0.16 | 0.01 ; 0.16 | 0.01 ; 0.16 | 0.01 ; 0.16 | 9 |
| CanH2 | # of adults co-feeding with a nymph (birds) | 0.01 ; 0.053 | 0.01 ; 0.053 | 0.01 ; 0.053 | 0.01 ; 0.053 | 0.01 ; 0.053 | 9 |
| ClaH2 | # of larvae co-feeding with an adult (birds) | 0.01 ; 0.22 | 0.01 ; 0.22 | 0.01 ; 0.22 | 0.01 ; 0.22 | 0.01 ; 0.22 | 9 |
| CnaH2 | # of nymphs co-feeding with an adult (birds) | 0.01 ; 0.024 | 0.01 ; 0.024 | 0.01 ; 0.024 | 0.01 ; 0.024 | 0.01 ; 0.024 | 9 |
| CaaH2 | # of adults co-feeding with an adult (birds) | 0.01 ; 0.014 | 0.01 ; 0.014 | 0.01 ; 0.014 | 0.01 ; 0.014 | 0.01 ; 0.014 | 9 |
| CllH3 | # of larvae co-feeding with a larva (primates) | 0.01 ; 1.06 | 0.01 ; 1.06 | 0.01 ; 1.06 | 0.01 ; 1.06 | 0.01 ; 1.06 | 10 |
| CnlH3 | # of nymphs co-feeding with a larva (primates) | 0.01; 0.30 | 0.01; 0.30 | 0.01; 0.30 | 0.01; 0.30 | 0.01; 0.30 | 10 |
| CalH3 | # of adults co-feeding with a larva (primates) | 0 ; 0 | 0 ; 0 | 0 ; 0 | 0 ; 0 | 0 ; 0 | 10 |
| ClnH3 | # of larvae co-feeding with a nymph (primates) | 0.01 ; 1.07 | 0.01 ; 1.07 | 0.01 ; 1.07 | 0.01 ; 1.07 | 0.01 ; 1.07 | 10 |
| CnnH3 | # of nymphs co-feeding with a nymph (primates) | 0.01 ; 0.38 | 0.01 ; 0.38 | 0.01 ; 0.38 | 0.01 ; 0.38 | 0.01 ; 0.38 | 10 |

| | | | | | | | |
|---|---|---|---|---|---|---|---|
| CanH3 | # of adults co-feeding with a nymph (primates) | 0 ; 0 | 0 ; 0 | 0 ; 0 | 0 ; 0 | 0 ; 0 | 10 |
| ClaH3 | # of larvae co-feeding with an adult (primates) | 0 ; 0 | 0 ; 0 | 0 ; 0 | 0 ; 0 | 0 ; 0 | 10 |
| CnaH3 | # of nymphs co-feeding with an adult (primates) | 0 ; 0 | 0 ; 0 | 0 ; 0 | 0 ; 0 | 0 ; 0 | 10 |
| CaaH3 | # of adults co-feeding with an adult (primates) | 0 ; 0 | 0 ; 0 | 0 ; 0 | 0 ; 0 | 0 ; 0 | 10 |
| CllH4 | # of larvae co-feeding with a larva (cattle) | 0 ; 0 | 0 ; 0 | 0 ; 0 | 0 ; 0 | 0 ; 0 | 11 |
| CnlH4 | # of nymphs co-feeding with a larva (cattle) | 0 ; 0 | 0 ; 0 | 0 ; 0 | 0 ; 0 | 0 ; 0 | 11 |
| CalH4 | # of adults co-feeding with a larva (cattle) | 0 ; 0 | 0 ; 0 | 0 ; 0 | 0 ; 0 | 0 ; 0 | 11 |
| ClnH4 | # of larvae co-feeding with a nymph (cattle) | 0 ; 0 | 0 ; 0 | 0 ; 0 | 0 ; 0 | 0 ; 0 | 11 |
| CnnH4 | # of nymphs co-feeding with a nymph (cattle) | 0.01 ; 0.1 | 0.01 ; 0.1 | 0.01 ; 0.1 | 0.01 ; 0.1 | 0.01 ; 0.1 | 11 |
| CanH4 | # of adults co-feeding with a nymph (cattle) | 0.01 ; 1.87 | 0.01 ; 1.87 | 0.01 ; 1.87 | 0.01 ; 1.87 | 0.01 ; 1.87 | 11 |
| ClaH4 | # of larvae co-feeding with an adult (cattle) | 0 ; 0 | 0 ; 0 | 0 ; 0 | 0 ; 0 | 0 ; 0 | 11 |
| CnaH4 | # of nymphs co-feeding with an adult (cattle) | 0.01 ; 0.4 | 0.01 ; 0.4 | 0.01 ; 0.4 | 0.01 ; 0.4 | 0.01 ; 0.4 | 11 |
| CaaH4 | # of adults co-feeding with an adult (cattle) | 0.01 ; 0.87 | 0.01 ; 0.87 | 0.01 ; 0.87 | 0.01 ; 0.87 | 0.01 ; 0.87 | 11 |
| NlhH1 | # of larvae on competent host (small mammals) | 0.01 ; 1.19 | 0.01 ; 1.19 | 0.01 ; 1.19 | 0.01 ; 1.19 | 0.01 ; 1.19 | 12 |
| NnhH1 | # of nymphs on competent host(small mammals) | 0.01 ; 2.63 | 0.01 ; 2.63 | 0.01 ; 2.63 | 0.01 ; 2.63 | 0.01 ; 2.63 | 11 |
| NahH1 | # of adults on competent host(small mammals) | 0 ; 0 | 0 ; 0 | 0 ; 0 | 0 ; 0 | 0 ; 0 | 11 |
| NlhH2 | # of larvae on competent host (birds) | 0.01 ; 10.58 | 0.01 ; 10.58 | 0.01 ; 10.58 | 0.01 ; 10.58 | 0.01 ; 10.58 | 8 |
| NnhH2 | # of nymphs on competent host(birds) | 0.01 ; 2.60 | 0.01 ; 2.60 | 0.01 ; 2.60 | 0.01 ; 2.60 | 0.01 ; 2.60 | 8 |
| NahH2 | # of adults on competent host(birds) | 0.01 ; 0.34 | 0.01 ; 0.34 | 0.01 ; 0.34 | 0.01 ; 0.34 | 0.01 ; 0.34 | 8 |
| NlhH3 | # of larvae on competent host(primates) | 0.01 ; 5.52 | 0.01 ; 5.52 | 0.01 ; 5.52 | 0.01 ; 5.52 | 0.01 ; 5.52 | 9 |
| NnhH3 | # of nymphs on competent host(primates) | 0.01 ; 1.58 | 0.01 ; 1.58 | 0.01 ; 1.58 | 0.01 ; 1.58 | 0.01 ; 1.58 | 9 |
| NahH3 | # of adults on competent host(primates) | 0 ; 0 | 0 ; 0 | 0 ; 0 | 0 ; 0 | 0 ; 0 | 9 |
| Dl | Days of attachment of larva | 2 ; 5 | 2 ; 5 | 2 ; 5 | 2 ; 5 | 2 ; 5 | 7 |
| Dn | Days of attachment of nymph | 7 ; 8 | 7 ; 8 | 7 ; 8 | 7 ; 8 | 7 ; 8 | 7 |
| Da | Days of attachment of adult | 10 ; 21 | 10 ; 21 | 10 ; 21 | 10 ; 21 | 10 ; 21 | 13 |

| | | | | | | | |
|---|---|---|---|---|---|---|---|
| IH1 | Systemic infection duration in host (small mammals) | 2 ; 7 | 2 ; 7 | 2 ; 7 | 2 ; 7 | 2 ; 7 | 7 |
| IH2 | Systemic infection duration in host (birds) | 2 ; 7 | 2 ; 7 | 2 ; 7 | 2 ; 7 | 2 ; 7 | 7 |
| IH3 | Systemic infection duration in host (primates) | 7 ; 12 | 7 ; 12 | 7 ; 12 | 7 ; 12 | 7 ; 12 | 7 |
| TT | Efficiency from tick to tick | 0.01 ; 1 | 0.01 ; 1 | 0.01 ; 1 | 0.01 ; 1 | 0.01 ; 1 | |
| Pl | Efficiency from competent host to larva | 0.6 ; 0.9 | 0.6 ; 0.9 | 0.6 ; 0.9 | 0.6 ; 0.9 | 0.6 ; 0.9 | 13,14 |
| Pn | Efficiency from competent host to nymph | 0.6 ; 0.9 | 0.6 ; 0.9 | 0.6 ; 0.9 | 0.6 ; 0.9 | 0.6 ; 0.9 | 13,14 |
| Pa | Efficiency from competent host to adult | 0.6 ; 0.9 | 0.6 ; 0.9 | 0.6 ; 0.9 | 0.6 ; 0.9 | 0.6 ; 0.9 | 13,14 |
| QlH1 | Efficiency from larva to competent host (small mammals) | 0.3 ; 0.75 | 0.3 ; 0.75 | 0.3 ; 0.75 | 0.3 ; 0.75 | 0.3 ; 0.75 | 13,15 |
| QnH1 | Efficiency from nymph to competent host (small mammals) | 0.3 ; 0.75 | 0.3 ; 0.75 | 0.3 ; 0.75 | 0.3 ; 0.75 | 0.3 ; 0.75 | 13,15 |
| QaH1 | Efficiency from adult to competent host (small mammals) | 0.3 ; 0.75 | 0.3 ; 0.75 | 0.3 ; 0.75 | 0.3 ; 0.75 | 0.3 ; 0.75 | 13,15 |
| QlH2 | Efficiency from larva to competent host (birds) | 0.3 ; 0.75 | 0.3 ; 0.75 | 0.3 ; 0.75 | 0.3 ; 0.75 | 0.3 ; 0.75 | 13,15 |
| QnH2 | Efficiency from nymph to competent host (birds) | 0.3 ; 0.75 | 0.3 ; 0.75 | 0.3 ; 0.75 | 0.3 ; 0.75 | 0.3 ; 0.75 | 13,15 |
| QaH2 | Efficiency from adult to competent host (birds) | 0.3 ; 0.75 | 0.3 ; 0.75 | 0.3 ; 0.75 | 0.3 ; 0.75 | 0.3 ; 0.75 | 13,15 |
| QlH3 | Efficiency from larva to competent host (primates) | 0.3 ; 0.75 | 0.3 ; 0.75 | 0.3 ; 0.75 | 0.3 ; 0.75 | 0.3 ; 0.75 | 13,15 |
| QnH3 | Efficiency from nymph to competent host (primates) | 0.3 ; 0.75 | 0.3 ; 0.75 | 0.3 ; 0.75 | 0.3 ; 0.75 | 0.3 ; 0.75 | 13,15 |
| QaH3 | Efficiency from adult to competent host (primates) | 0.3 ; 0.75 | 0.3 ; 0.75 | 0.3 ; 0.75 | 0.3 ; 0.75 | 0.3 ; 0.75 | 13,15 |
| Ra | Efficiency from adult to egg | 0.01 ; 0.1 | 0.01 ; 0.1 | 0.01 ; 0.1 | 0.01 ; 0.1 | 0.01 ; 0.1 | 7 |
| Hc | Fraction of blood meals on hosts competent for systemic and non-systemic transmission | 0.5 ; 0.9 | 0.5 ; 0.9 | 0.5 ; 0.9 | 0.5 ; 0.9 | 0.5 ; 0.9 | |
| Cs | Fraction of ticks feeding close enough for non-systemic transmission | 0.1 ; 0.5 | 0.1 ; 0.5 | 0.1 ; 0.5 | 0.1 ; 0.5 | 0.1 ; 0.5 | 7 |

### S1.2 Parameters for used scaling factors

Below are the parameters used for the scaling factors. Note that the parameter range for eggs varies in the cattle density model. The minimum of the range is used as eggs are increasing with cattle density and the specified range ensures that the number of eggs does not exceed 3526 in the model.

| Parameter | Description | Evergreen forest | Deciduous forest | Plantation | Paddy | Human habitation | Reference |
|---|---|---|---|---|---|---|---|
| Np | 1- proportion decrease in ticks | 1 ; 1 | 1 ; 1 | 0.56 ; 0.56 | 0.56 ; 0.56 | 0.69 ; 0.69 | |
| E (habitat type) | Mean # of eggs per adult | 1090;3536 | 1090;3536 | 1090;3536 | 1090;3536 | 1090;3536 | 16 |
| rL | proportion increase in tick abundance and burden with increase 1 cattle per km2 | 0.020 ; 0.032 | 0.020 ; 0.032 | 0.020 ; 0.032 | 0.020 ; 0.032 | 0.020 ; 0.032 | 17 |
| LD | cattle density (ind per km2) | 40.1 ; 40.1 | 60 ; 60 | 60 ; 60 | 44.64 ; 44.64 | 70 ; 70 | 8 |
| E (cattle density) | Mean # of eggs per adult | 1090;1090 | 1090;1090 | 1090;1090 | 1090;1090 | 1090;1090 | 16 |

### S1.3 Equations used in NGM

Below are the equations used for each matrix element in the 7x7 matrix. Note that when transovarial transmission is excluded the first column and first row are removed from the matrix. The equations outline the model using cattle density as a scaling factor using $(L_d r_L + 1)$, this term is replaced with $N_p$ when using habitat type as a scaling factor.

$$K = \begin{bmatrix} k_{1,1} & k_{1,2} & k_{1,3} & k_{1,4} & k_{1,5} & k_{1,6} & k_{1,7} \\ k_{2,1} & k_{2,2} & k_{2,3} & k_{2,4} & k_{2,5} & k_{2,6} & k_{2,7} \\ k_{3,1} & k_{3,2} & k_{3,3} & k_{3,4} & k_{3,5} & k_{3,6} & k_{3,7} \\ k_{4,1} & k_{4,2} & k_{4,3} & k_{4,4} & k_{4,5} & k_{4,6} & k_{4,7} \\ k_{5,1} & k_{5,2} & k_{5,3} & k_{5,4} & k_{5,5} & k_{5,6} & k_{5,7} \\ k_{6,1} & k_{6,2} & k_{6,3} & k_{6,4} & k_{6,5} & k_{6,6} & k_{6,7} \\ k_{7,1} & k_{7,2} & k_{7,3} & k_{7,4} & k_{7,5} & k_{7,6} & k_{7,7} \end{bmatrix}$$

Transovarial transmission

***Number of ticks infected as an egg by one tick infected as…***

***an egg***

$$k_{1,1} = (L_d r_L + 1) E S_l S_n S_a R_a$$

***a larva***

$$k_{1,2} = (L_d r_L + 1) E S_n S_a R_a$$

***a nymph***

$$k_{1,3} = (L_d r_L + 1) E S_a R_a$$

***an adult***

$$k_{1,4} = (L_d r_L + 1) E R_a$$

Non-systemic transmission

$$sm = small\ mammals, bir = birds, pri = primates\ and\ cat = cattle$$

***Number of larvae infected by one tick infected as…***

***an egg***

$k_{2,1} = S_l H_c\big((L_d r_L + 1)Cll_{cat} C_s \theta_{TT} \theta Pb_{cat} + (L_d r_L + 1)Cll_{pri} C_s \theta_{TT} Pb_{pri} + (L_d r_L + 1)Cll_{bir} C_s \theta_{TT} Pb_{bir} + (L_d r_L + 1)Cll_{sm} C_s \theta_{TT} Pb_{sm}\big) + S_l S_n H_c\big((L_d r_L + 1)Cln_{cat} C_s \theta_{TT} Pb_{cat} + (L_d r_L + 1)Cln_{pri} C_s \theta_{TT} Pb_{pri} + (L_d r_L + 1)Cln_{bir} C_s \theta_{TT} Pb_{bir} + (L_d r_L + 1)Cln_{sm} C_s \theta_{TT} Pb_{sm}\big) + S_l S_n S_a H_c\big((L_d r_L + 1)Cla_{cat} C_s \theta_{TT} Pb_{cat} + (L_d r_L + 1)Cla_{pri} C_s \theta_{TT} Pb_{pri} + (L_d r_L + 1)Cla_{bir} C_s \theta_{TT} Pb_{bir} + (L_d r_L + 1)Cla_{sm} C_s \theta_{TT} Pb_{sm}\big)$

***a larva***

$k_{2,2} = S_n H_c\big((L_d r_L + 1)Cln_{cat} C_s \theta_{TT} Pb_{cat} + (L_d r_L + 1)Cln_{pri} C_s \theta_{TT} Pb_{pri} + (L_d r_L + 1)Cln_{bir} C_s \theta_{TT} Pb_{bir} + (L_d r_L + 1)Cln_{sm} C_s \theta_{TT} Pb_{sm}\big) + S_n S_a H_c\big((L_d r_L + 1)Cla_{cat} C_s \theta_{TT} Pb_{cat} + (L_d r_L + 1)Cla_{pri} C_s \theta_{TT} Pb_{pri} + (L_d r_L + 1)Cla_{bir} C_s \theta_{TT} Pb_{bir} + (L_d r_L + 1)Cla_{sm} C_s \theta_{TT} Pb_{sm}\big)$

***a nymph***

$k_{2,3} = S_a H_c\big((L_d r_L + 1)Cla_{cat} C_s \theta_{TT} Pb_{cat} + (L_d r_L + 1)Cla_{pri} C_s \theta_{TT} Pb_{pri} + (L_d r_L + 1)Cla_{bir} C_s \theta_{TT} Pb_{bir} + (L_d r_L + 1)Cla_{sm} C_s \theta_{TT} Pb_{sm}\big)$

***Number of nymphs infected by one tick infected as…***

***an egg***

$k_{3,1} = S_l H_c\big((L_d r_L + 1)Cnl_{cat} C_s \theta_{TT} \theta Pb_{cat} + (L_d r_L + 1)Cnl_{pri} C_s \theta_{TT} Pb_{pri} + (L_d r_L + 1)Cnl_{bir} C_s \theta_{TT} Pb_{bir} + L_d r_L Cnl_{sm} C_s \theta_{TT} Pb_{sm}\big) + S_l S_n H_c\big((L_d r_L + 1)Cnn_{cat} C_s \theta_{TT} Pb_{cat} + (L_d r_L + 1)Cnn_{pri} C_s \theta_{TT} Pb_{pri} + (L_d r_L + 1)Cnn_{bir} C_s \theta_{TT} Pb_{bir} + (L_d r_L + 1)Cnn_{sm} C_s \theta_{TT} Pb_{sm}\big) + S_l S_n S_a H_c\big((L_d r_L + 1)Cna_{cat} C_s \theta_{TT} Pb_{cat} + (L_d r_L + 1)Cna_{pri} C_s \theta_{TT} Pb_{pri} + (L_d r_L + 1)Cna_{bir} C_s \theta_{TT} Pb_{bir} + (L_d r_L + 1)Cna_{sm} C_s \theta_{TT} Pb_{sm}\big)$

***a larva***

$k_{3,2} = S_n H_c\big((L_d r_L + 1)Cnn_{cat} C_s \theta_{TT} Pb_{cat} + (L_d r_L + 1)Cnn_{pri} C_s \theta_{TT} Pb_{pri} + (L_d r_L + 1)Cnn_{bir} C_s \theta_{TT} Pb_{bir} + (L_d r_L + 1)Cnn_{sm} C_s \theta_{TT} Pb_{sm}\big) + S_n S_a H_c\big((L_d r_L + 1)Cna_{cat} C_s \theta_{TT} Pb_{cat} + (L_d r_L + 1)Cna_{pri} C_s \theta_{TT} Pb_{pri} + (L_d r_L + 1)Cna_{bir} C_s \theta_{TT} Pb_{bir} + (L_d r_L + 1)Cna_{sm} C_s \theta_{TT} Pb_{sm}\big)$

***a nymph***

$$k_{3,3} = S_a H_c\big((L_d r_L + 1)Cna_{cat} C_s \theta_{TT} Pb_{cat} + (L_d r_L + 1)Cna_{pri} C_s \theta_{TT} Pb_{pri} + (L_d r_L + 1)Cna_{bir} C_s \theta_{TT} Pb_{bir} + (L_d r_L + 1)Cna_{sm} C_s \theta_{TT} Pb_{sm}\big)$$

***Number of adults infected by one tick infected as…***

***an egg***

$$k_{4,1} = S_l H_c\big((L_d r_L + 1)Cal_{cat} C_s \theta_{TT} \theta Pb_{cat} + (L_d r_L + 1)Cal_{pri} C_s \theta_{TT} Pb_{pri} + (L_d r_L + 1)Cal_{bir} C_s \theta_{TT} Pb_{bir} + (L_d r_L + 1)Cal_{sm} C_s \theta_{TT} Pb_{sm}\big) + S_l S_n H_c\big((L_d r_L + 1)Can_{cat} C_s \theta_{TT} Pb_{cat} + (L_d r_L + 1)Can_{pri} C_s \theta_{TT} Pb_{pri} + (L_d r_L + 1)Can_{bir} C_s \theta_{TT} Pb_{bir} + (L_d r_L + 1)Can_{sm} C_s \theta_{TT} Pb_{sm}\big) + S_l S_n S_a H_c\big((L_d r_L + 1)Caa_{cat} C_s \theta_{TT} Pb_{cat} + (L_d r_L + 1)Caa_{pri} C_s \theta_{TT} Pb_{pri} + (L_d r_L + 1)Caa_{bir} C_s \theta_{TT} Pb_{bir} + (L_d r_L + 1)Caa_{sm} C_s \theta_{TT} Pb_{sm}\big)$$

***a larva***

$$k_{4,2} = S_n H_c\big((L_d r_L + 1)Can_{cat} C_s \theta_{TT} Pb_{cat} + (L_d r_L + 1)Can_{pri} C_s \theta_{TT} Pb_{pri} + (L_d r_L + 1)Can_{bir} C_s \theta_{TT} Pb_{bir} + (L_d r_L + 1)Can_{sm} C_s \theta_{TT} Pb_{sm}\big) + S_n S_a H_c\big((L_d r_L + 1)Caa_{cat} C_s \theta_{TT} Pb_{cat} + (L_d r_L + 1)Caa_{pri} C_s \theta_{TT} Pb_{pri} + (L_d r_L + 1)Caa_{bir} C_s \theta_{TT} Pb_{bir} + (L_d r_L + 1)Caa_{sm} C_s \theta_{TT} Pb_{sm}\big)$$

***a nymph***

$$k_{4,3} = S_a H_c\big((L_d r_L + 1)Caa_{cat} C_s \theta_{TT} Pb_{cat} + (L_d r_L + 1)Caa_{pri} C_s \theta_{TT} Pb_{pri} + (L_d r_L + 1)Caa_{bir} C_s \theta_{TT} Pb_{bir} + (L_d r_L + 1)Caa_{sm} C_s \theta_{TT} Pb_{sm}\big)$$

<u>Systemic transmission</u>

$$sm = small\ mammals, bir = birds, pri = primates\ and\ cat = cattle$$

***Number of larvae infected by an infected…***

***small mammal***

$$k_{2,5} = \frac{(L_d r_L + 1)Nl_{sm} Ih_{sm} P_l}{D_l}$$

***bird***

$$k_{2,6} = \frac{(L_d r_L + 1)Nl_{bir} Ih_{bir} P_l}{D_l}$$

***primate***

$$k_{2,7} = \frac{(L_d r_L + 1)Nl_{pri} Ih_{pri} P_l}{D_l}$$

***Number of nymphs infected by an infected…***

***small mammal***

$$k_{3,5} = \frac{(L_d r_L + 1) N n_{sm} I h_{sm} P_n}{D_n}$$

***bird***

$$k_{3,6} = \frac{(L_d r_L + 1) N n_{bir} I h_{bir} P_n}{D_n}$$

***primate***

$$k_{3,7} = \frac{(L_d r_L + 1) N n_{pri} I h_{pri} P_n}{D_n}$$

***Number of adults infected by an infected…***

***small mammal***

$$k_{4,5} = \frac{(L_d r_L + 1) N a_{sm} I h_{sm} P_a}{D_a}$$

***bird***

$$k_{4,6} = \frac{(L_d r_L + 1) N a_{bir} I h_{bir} P_a}{D_a}$$

***primate***

$$k_{4,7} = \frac{(L_d r_L + 1) N a_{pri} I h_{pri} P_a}{D_a}$$

***Number of small mammals infected by a tick infected as …***

***an egg***

$$k_{5,1} = (S_l Q l_{sm} H_c + S_l S_n Q n_{sm} H_c + S_l S_n S_a Q a_{sm} H_c)\ P b_{sm}$$

***a larva***

$$k_{5,2} = (S_n Q n_{sm} H_c + S_n S_a Q a_{sm} H_c)\ P b_{sm}$$

**a nymph**

$$k_{5,3} = (S_a Q a_{sm} H_c)\ Pb_{sm}$$

***Number of birds infected by a tick infected as ..***

***an egg***

$$k_{6,1} = (S_l Q l_{bir} H_c + S_l S_n Q n_{bir} H_c + S_l S_n S_a Q a_{bir} H_c)\ Pb_{bir}$$

**a larva**

$$k_{6,2} = (S_n Q n_{bir} H_c + S_n S_a Q a_{bir} H_c)\ Pb_{bir}$$

**a nymph**

$$k_{6,3} = (S_a Q a_{bir} H_c)\ Pb_{bir}$$

***Number of primates infected by a tick infected as ..***

***an egg***

$$k_{7,1} = (S_l Q l_{pri} H_c + S_l S_n Q n_{pri} H_c + S_l S_n S_a Q a_{pri} H_c)\ Pb_{pri}$$

***a larva***

$$k_{7,2} = (S_n Q n_{pri} H_c + S_n S_a Q a_{pri} H_c)\ Pb_{pri}$$

***a nymph***

$$k_{7,3} = (S_a Q a_{pri} H_c)\ Pb_{pri}$$

### S1.4 Bird species used to estimate bird densities

| Common name | Scientific name |
| --- | --- |

| | |
|---|---|
| Blyth's Reed Warbler | *Acrocephalus dumetorum* |
| Indian Blackbird | *Turdus simillimus* |
| Gray Junglefowl | *Gallus sonneratii* |
| Greater Coucal | *Centropus sinensis* |
| Indian Bushlark | *Mirafra erythroptera* |
| Indian Pitta | *Pitta brachyura* |
| Jungle Babbler | *Turdoides striata* |
| Jungle Myna | *Acridotheres fuscus* |
| Long-tailed Shrike | *Lanius schach* |
| Orange-headed Thrush | *Geokichla citrina* |
| Oriental Magpie-Robin | *Copsychus saularis* |
| Puff-throated Babbler | *Pellorneum ruficeps* |

| | |
|---|---|
| Rufous Babbler | *Turdoides subrufa* |
| Shikra | *Accipiter badius* |
| White-bellied Drongo | *Dicrurus caerulescens* |
| Yellow-billed Babbler | *Turdoides affinis* |
