## Supplementary Information S2 for "Using mechanistic models to highlight research priorities for tick-borne zoonotic diseases: Improving our understanding of the ecology and maintenance of Kyasanur Forest Disease in India"

### Supplementary S2: Additional results from models.

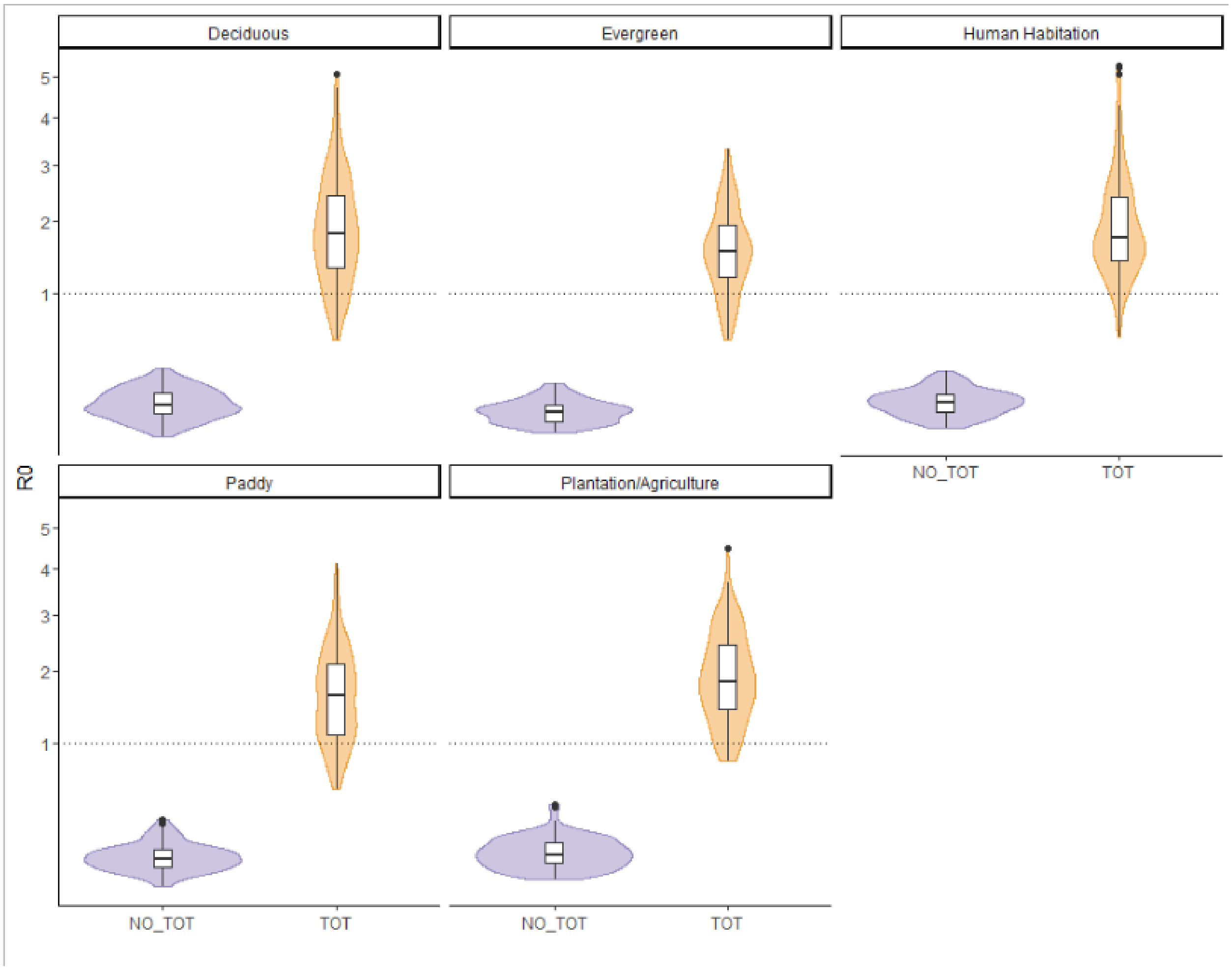


Figure S2.1: Distribution of R0 values in each habitat type for models using scaling factor based on cattle density. Plot shows results for models excluding transovarial transmission (NO_TOT) and including transovarial transmission (TOT) in different habitat types. R0 is significantly lower in all scenarios when TOT is excluded from the model.

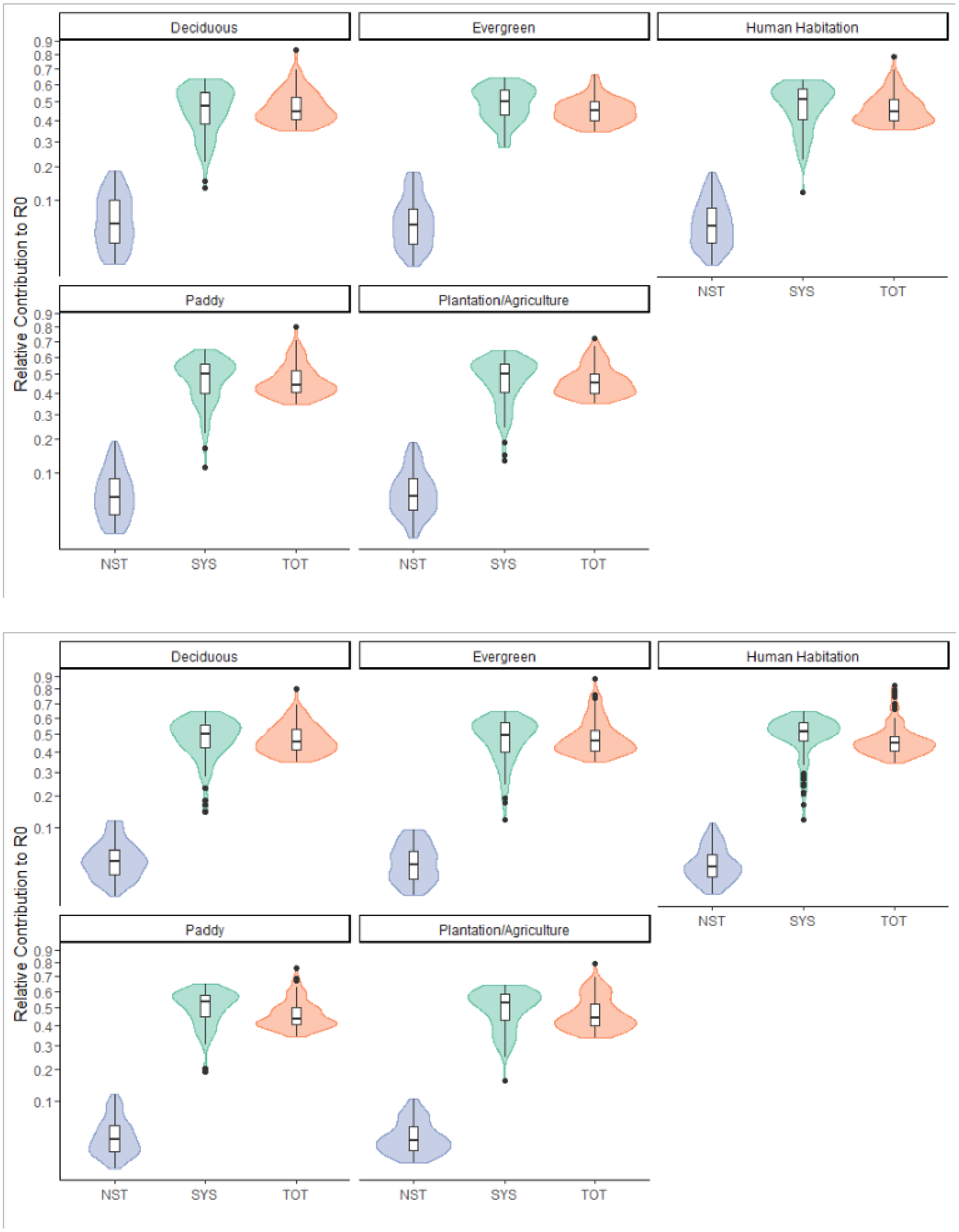


Figure S.2.2: Distribution of values for relative contribution to R0 for each route of transmission with transovarial transmission included in the model (top) using cattle density as a scaling factor and (bottom) using habitat type as a scaling factor. Grid shows results for each habitat type. Across all scenarios both systemic and transovarial had an equal contribution to R0, which was significantly higher than non-systemic transmission.

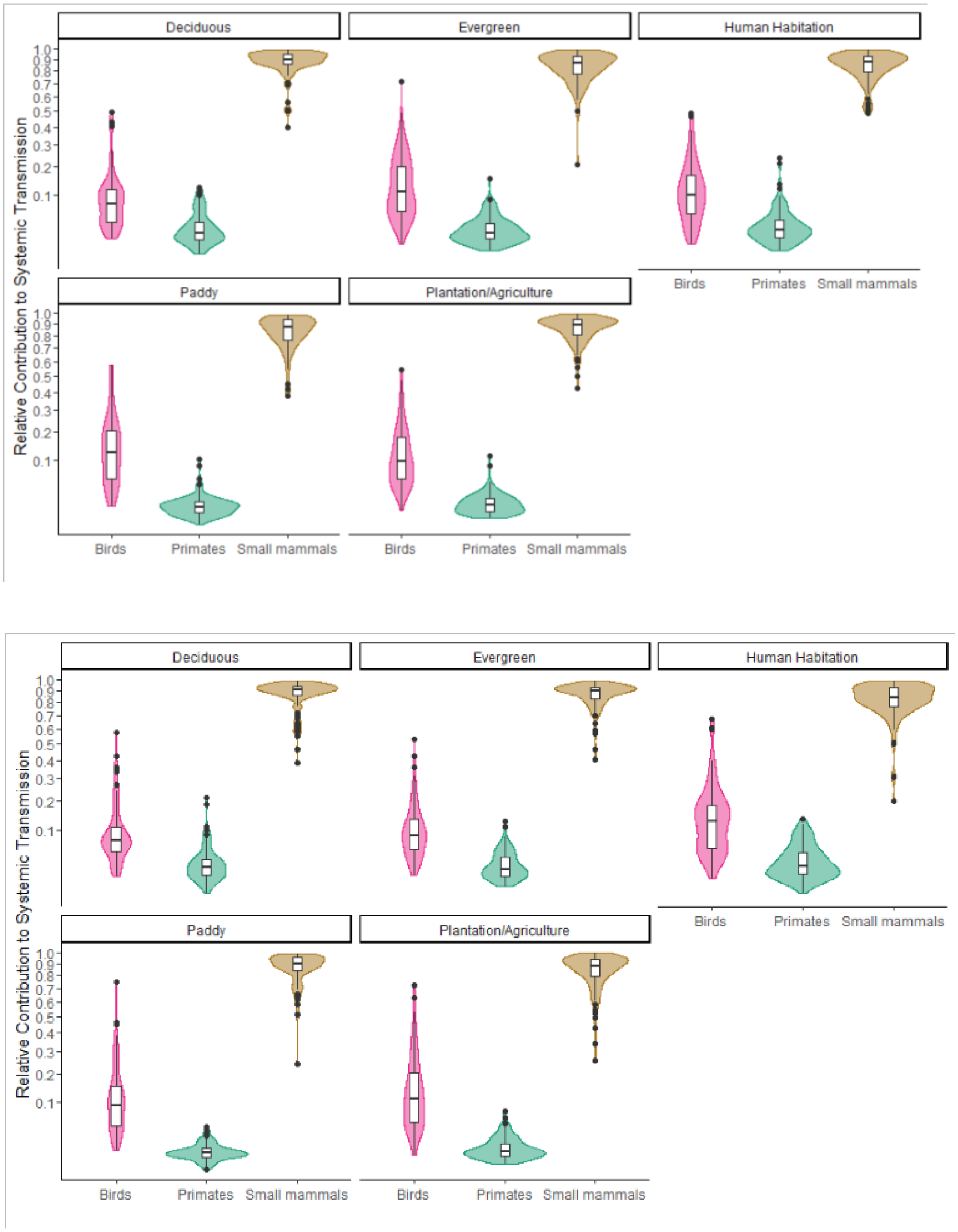


Figure S.2.3: Distribution of values for relative contribution to systemic transmission for each host with transovarial transmission included in the model (top) using cattle density as a scaling factor and (bottom) using habitat type as a scaling factor. Grid shows results for each habitat type.

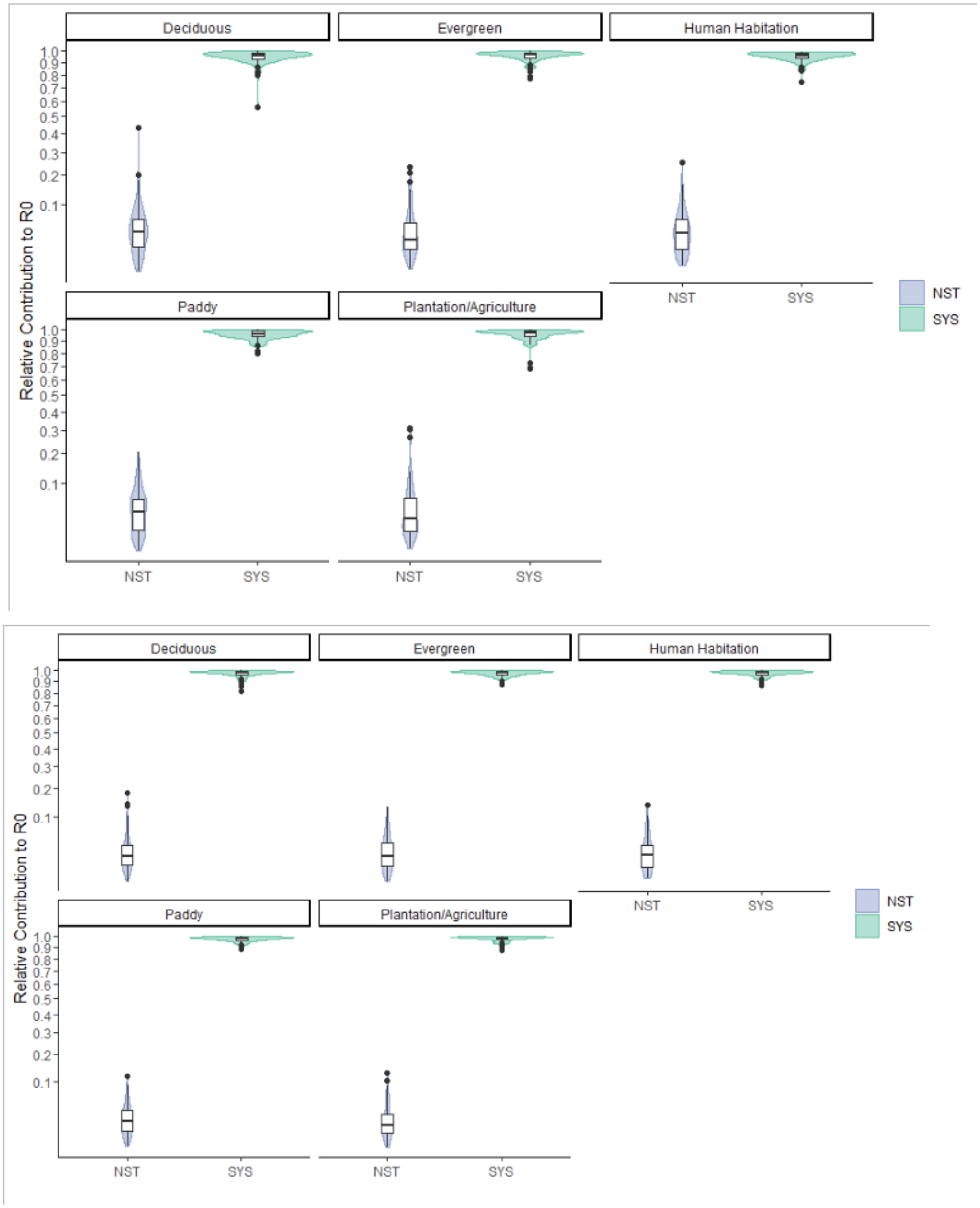


Figure S.2.4: Distribution of values for relative contribution to R0 for each route of transmission with transovarial transmission excluded from the model (top) using cattle density as a scaling factor and (bottom) using habitat type as a scaling factor. Grid shows results for each habitat type. Without transovarial transmission included in the model, systemic transmission had the highest contribution to R0 and non-systemic transmission had the lowest contribution.

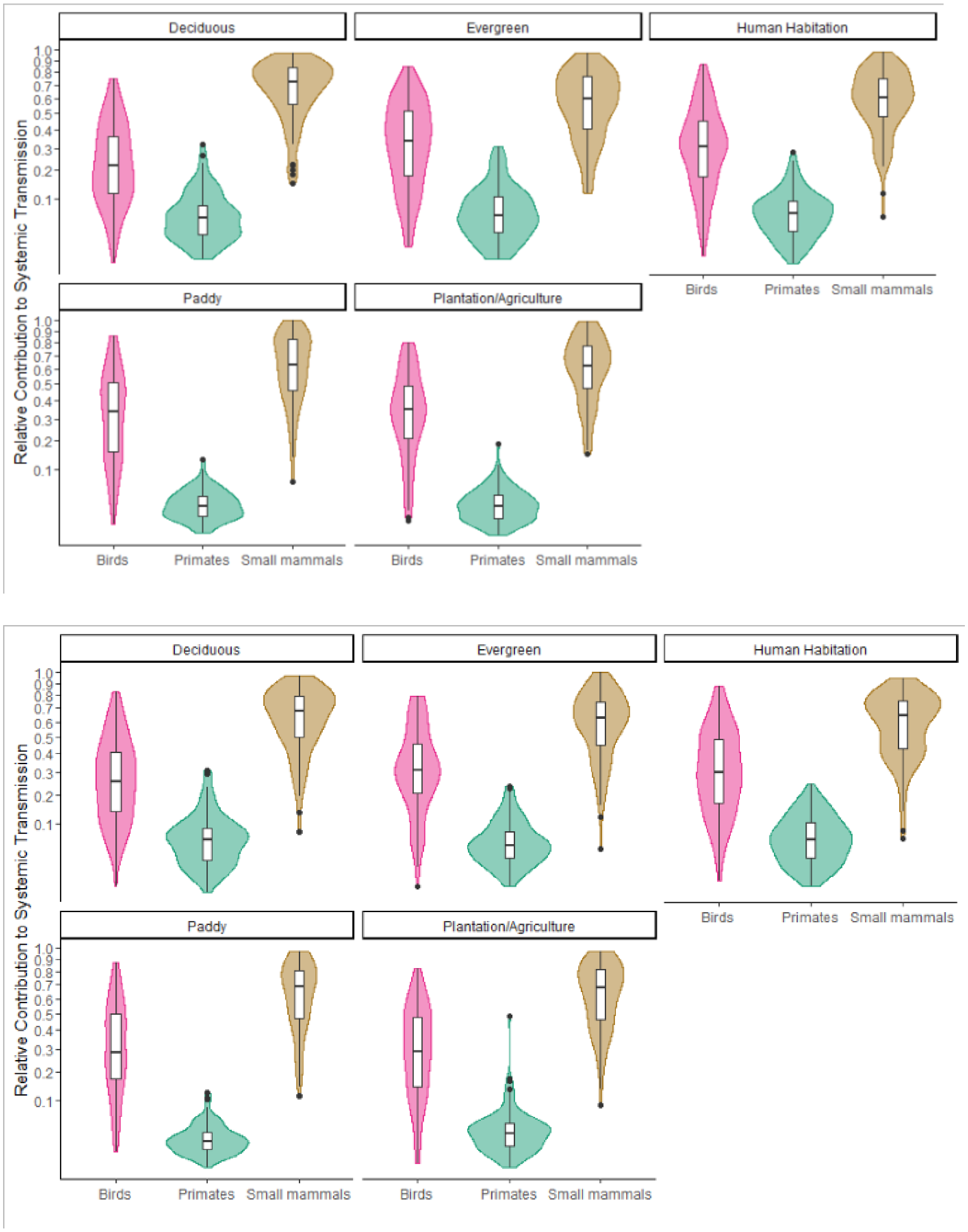


Figure S.2.3: Distribution of values for relative contribution to systemic transmission for each host with transovarial transmission excluded from the model (top) using cattle density as a scaling factor and (bottom) using habitat type as a scaling factor. Grid shows results for each habitat type. Across all scenarios, small mammals were predicted to have the greatest contribution to systemic transmission. This was follow by birds and then finally primates which had the lowest contribution to systemic transmission.
